# An Essential Role of UBXN3B in B Lymphopoiesis

**DOI:** 10.1101/2021.03.04.433919

**Authors:** Tingting Geng, Duomeng Yang, Tao Lin, Andrew G. Harrison, Binsheng Wang, Blake Torrance, Kepeng Wang, Yanlin Wang, Long Yang, Laura Haynes, Gong Cheng, Anthony T. Vella, Erol Fikrig, Penghua Wang

**Affiliations:** Department of Immunology, School of Medicine, UConn Health, Farmington, CT 06030, USA; Center on Aging and Department of Genetics and Genome Sciences, School of Medicine, UConn Health, Farmington, CT 06030, USA; Department of Medicine, School of Medicine, UConn Health, Farmington, CT 06030, USA; School of integrative Medicine, Tianjin University of Traditional Chinese Medicine, Tianjin 301617, China; Department of Basic Sciences, School of Medicine, Tsinghua University, Beijing, China; Section of Infectious Diseases, Yale University School of Medicine, 333 Cedar Street, New Haven, CT, 06510, USA

**Keywords:** UBXN, FAF2, hematopoiesis, B lymphopoiesis, SARS-CoV-2

## Abstract

Hematopoiesis is finely regulated to enable timely production of the right numbers and types of mature immune cells to maintain tissue homeostasis. Dysregulated hematopoiesis may compromise antiviral immunity and/or exacerbate immunopathogenesis. Herein, we report an essential role of UBXN3B in maintenance of hematopoietic homeostasis and restriction of immunopathogenesis during respiratory viral infection. *Ubxn3b* deficient (*Ubxn3b*^−/−^) mice are highly vulnerable to SARS-CoV-2 and influenza A infection, characterized by more severe lung immunopathology, lower virus-specific IgG, significantly fewer B cells, but more myeloid cells than *Ubxn3b*^+/+^ littermates. This aberrant immune compartmentalization is recapitulated in uninfected *Ubxn3b*^−/−^ mice. Mechanistically, UBXN3B controls precursor B-I (pre-BI) transition to pre-BII and subsequent proliferation in a cell-intrinsic manner, by maintaining BLNK protein stability and pre-BCR signaling. These results reveal an essential role of UBXN3B for the early stage of B cell development.

## INTRODUCTION

The immune system is comprised of various cell types that coordinate responses to infection, maintain tissue and immune homeostasis. Peripheral immune cells, with the exception of a few cell types such as long-lived memory T cells and some tissue macrophages, are constantly replenished from bone marrow stem cells through progenitor cells ^1^. For instance, approximately 0.5–1 × 10^11^ granulocytes are generated daily in adult human individuals ^2^. The hematopoietic system is a hierarchically organized, somatic stem cell-maintained organ system, with long-lived and self-renewing pluripotent hematopoietic stem cells (LT-HSCs) at its apex ^1^. LT-HSCs differentiate into short-term multipotent progenitors (MPPs or ST-HSCs) and lineage-committed hematopoietic progenitors, which in turn will eventually differentiate into the numerous mature blood cell lineages ^3^. While at the apex of the hematopoietic hierarchy, LT-HSCs are largely quiescent, and the highly proliferative MPPs are the primary contributor to steady-state hematopoiesis ^4 5^. MPPs are capable of differentiating into lineage-committed progenitors, e.g., common lymphoid progenitors (CLPs) and common myeloid progenitors (CMP), which turn into blast cells leading to specific cell types. Among hematopoiesis, B cell development is the best studied, with several stages clearly defined, including pre-progenitor (pre-pro) B, precursor B I (pre-BI), large pre-BII, small pre-BII and immature B (imm-B) inside bone marrow. In bone marrow, pre-BI transition to large pre-BII is considered an essential checkpoint, involving rearrangement of variable (V)/diversity (D)/joining (J) gene segments by recombination activating genes (RAG1/2) and assembly of pre-B cell receptor (pre-BCR) with a surrogate light chain (SLC) and immunoglobulin (Ig) μ heavy chain (μH) on cell surface. Pre-BI cells without a functional pre-BCR undergo apoptosis. Once passing the first quality check, pre-BI becomes large pre-BII, which proliferates several rounds and turn into small pre-BII. At this stage, small pre-BII no longer expresses SLC, but begins expressing an Ig light chain κ or λ that forms a BCR together with μH, and becomes imm-B (checkpoint 2). Imm-B cells exit bone marrow and mature in the peripheral immune organs such as spleen and lymph node ^6^. This process is controlled by a unique set of cell-intrinsic transcription factors and cell-extrinsic factors such as cytokines, chemokines and growth factors in its bone marrow niche ^3^.

The human genome encodes 13 ubiquitin regulatory X (UBX) domain-containing proteins, designated UBXNs. The UBX domain shares weak homology with ubiquitin but adopts the same three dimensional fold as ubiquitin ^7^. Many UBXNs are capable of binding multiple E3 ubiquitin ligases and p97 (also known as VCP), an ATPase associated with various cellular activities (AAA ATPase) ^8,9^ However, the physiological functions of UBXNs remain poorly characterized. We and other research groups have recently shown that several UBXNs regulate viral RNA-sensing RIG-I (retinoic acid inducible gene 1) like receptor-mitochondrial antiviral viral signaling (RLR-MAVS) ^10–12^ and Nuclear factor-κB (NF-κB) signaling pathways ^13,14^. Of note, we recently reported that UBXN3B controls a DNA virus infection by positively regulating the dsDNA-sensing cGAS (cyclic di-GMP-AMP synthase)-STING (stimulator-of-interferon-genes) signaling and innate immunity ^15^. However, the physiological function of UBXN3B in RNA virus infection remains unknown. To this end, we will study two important respiratory viruses, including one positive-sense single-stranded RNA [(+) ssRNA] severe acute respiratory syndrome (COVID-19)-causing coronavirus 2 (SARS-CoV-2), and a negative-sense single-stranded RNA [(-) ssRNA] influenza A virus (IAV). Both viruses induce life-threatening lung immunopathology, typified by elevated levels of inflammatory mediators, myeloid immune infiltrates in the lung, neutrophilia and lymphopenia^16 17^. Herein, we report that *Ubxn3b* deficient (*Ubxn3b*^−/−^) mice are highly vulnerable to SARS-CoV-2 and IAV infection, typified by higher viral loads and inflammatory mediators, more severe immunopathology in the lung, but lower virus-specific immunoglobulin (Ig) G and slower resolution of disease, when compared to *Ubxn3*^+/+^ littermates. Of note, SARS-CoV-2 infected *Ubxn3b*^−/−^ mice have lower B/T cell counts, while more myeloid cells, and consequently a higher neutrophil-to-lymphocyte ratio (N/L) in the lung and blood. Intriguingly, this abnormal immune compartmentalization is also recapitulated in uninfected *Ubxn3b*^−/−^ mice when compared to *Ubxn3b*^+/+^ littermates. Reciprocal bone marrow transplantation reveals that the B cell defect in *Ubxn3b*^−/−^ is cell-intrinsic. Mechanistically, UBXN3B controls precursor B-I (pre-BI) transition to pre-BII and subsequent proliferation in a cell-intrinsic manner, by maintaining BLNK protein stability and pre-BCR signaling.

## RESULTS

### UBXN3B restricts pathogenesis of respiratory viruses

We have long been interested in UBXNs because of their potential function in ubiquitination and immune regulation. Using a tamoxifen-inducible Cre-LoxP system, we recently successfully deleted an essential gene, UBXN3B, in adult mice and demonstrated that UBXN3B positively regulates the STING-mediated type I IFN response to a DNA virus^15^. Because STING signaling also plays a role in controlling some RNA viruses in an IFN-dependent or -independent manner, we continued investigating the physiological role of UBXN3B in restricting RNA virus infection. To this end, we tested with respiratory viruses of public health significance including SARS-CoV-2 and influenza A virus (IAV). Because mice are barely permissive to clinical isolates of SARS-CoV-2, we delivered human angiotensin-converting enzyme 2 (ACE2, the cellular entry receptor for SARS-CoV)-expressing Ad5 vector (replication-defective adenovirus vector) intranasally to the mouse lung before infection ^18 19^. We observed a slight drop in body mass a few days post SARS-CoV-2 infection (p.i.) and rapid recovery of *Ubxn3b*^+/+^ (Cre^+^ *Ubxn3b* ^flox/flox^ treated with corn oil) mice, while a ~10% reduction in the body weight of *Ubxn3b*^−/−^ (Cre^+^ *Ubxn3b* ^flox/flox^ treated with tamoxifen dissolved in corn oil) littermates by days 3-4 p.i. and a significant delay in recovery (**Fig.1a**). Moreover, all infected *Ubxn3b*^−/−^ mice showed hunched posture and decreased mobility at day 2 p.i., while *Ubxn3b*^+/+^ animals behaved normally (Suppl Movie 1 and 2). Histopathological analyses by hematoxylin and eosin (H&E) staining revealed more immune infiltrates in the lung of both *Ubxn3b*^+/+^ and *Ubxn3b*^−/−^ mice at day 3 p.i., compared to uninfected mice (day 0). However, there was no significant difference in the numbers of immune infiltrates between the two groups (**Fig.1b**). At day 10 p.i., many clusters of brownish cells were noted in the H&E sections of all *Ubxn3b*^−/−^, but none in any *Ubxn3b*^+/+^ mice (**Fig.1b**). We reasoned that these brownish cells were representative of hemosiderosis, a form of iron overload disorder resulting in the accumulation of hemosiderin, an iron-storage complex. In the lung, macrophages phagocytose red blood cells due to vascular leakage, leading to iron overload. Using iron staining we tested this hypothesis and detected a few lightly iron-laden cells at day 3 p.i., but many clusters of heavily iron-laden cells by day 10 p.i. in all the *Ubxn3b*^−/−^ lungs compared to *Ubxn3b*^+/+^ (**Fig.1c, d**). On day 35 p.i., we still noted moderate lung hemosiderosis in some knockout mice. Of note, with either low or high viral loads, all *Ubxn3b*^−/−^ mice presented a similar degree of hemosiderosis at days 3 and 10 p.i., while no *Ubxn3b*^+/+^ mouse had it (**Fig. 1c, d**), suggesting that the severe lung damage in *Ubxn3b*^−/−^ mice is primarily caused by immunopathology. We next asked if these observations could be extended to other respiratory viruses, such as influenza. Indeed, *Ubxn3b*^−/−^ mice lost more body weight; and of note, 70% of them succumbed to a dose of H1N1 influenza A that was only sublethal to *Ubxn3b*^+/+^ mice (**Fig.1e, f**).

**Fig.1.**
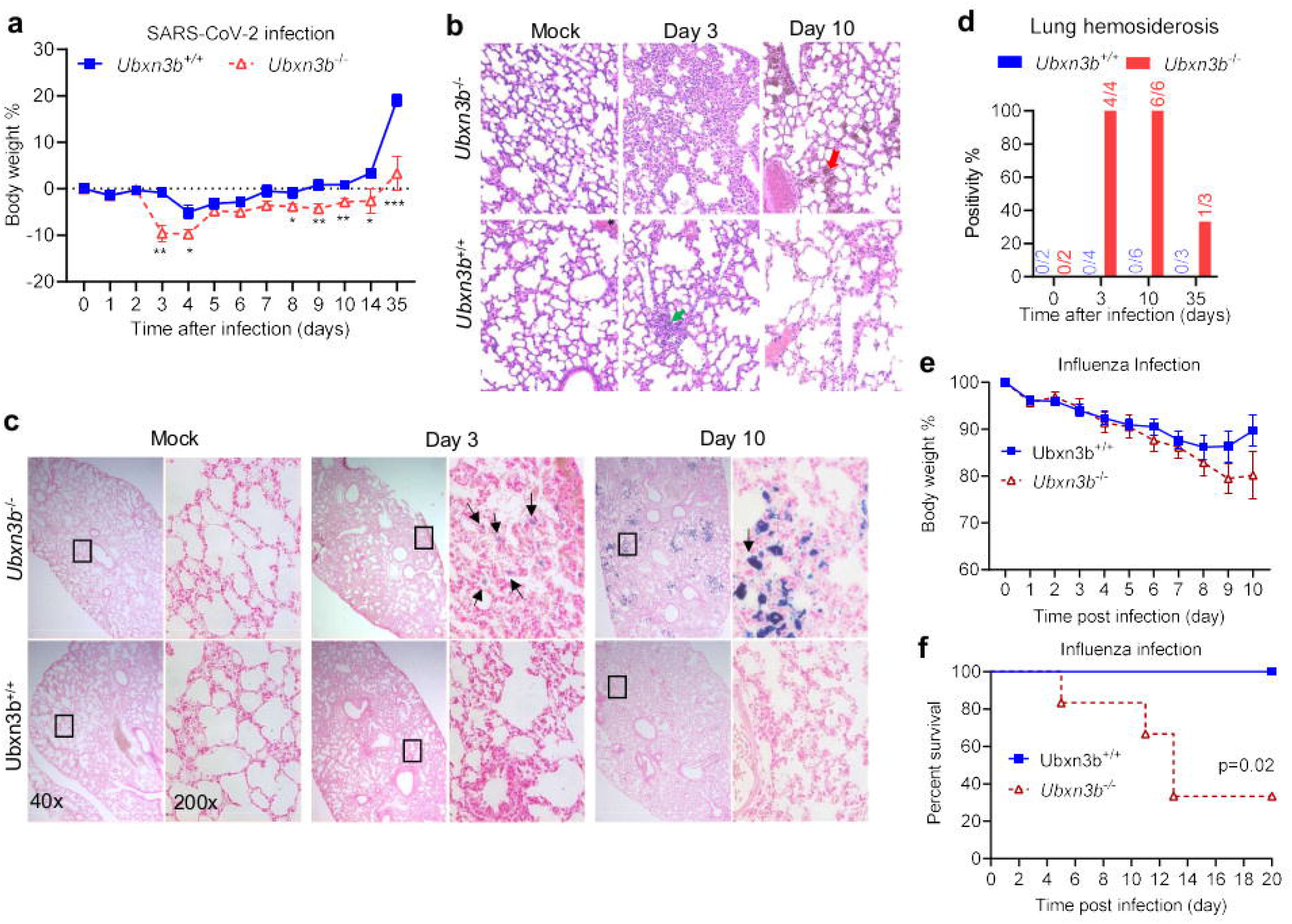
UBXN3B is essential for controlling SARS-CoV-2 and influenza pathogenesis. **a-d**) Sex-and-age matched mice were administered 2×10^5^ plaque forming units (PFU) of SARS-CoV-2 intranasally. **a**) Percentage changes in the body mass of mock-treated Cre^+^ Ubxn3b^flox/flox^ (designated *Ubxn3b*^+/+^) and tamoxifen (TMX) -treated Cre^+^ Ubxn3b^flox/flox^ (designated *Ubxn3b*^−/−^) littermates, during the course of SARS-CoV-2 infection. Data point: mean ± s.e.m, N=6-8. *, p<0.05; **, p<0.01; ***, p<0.001 (two-tailed Student’s *t*-test). **b**) Representative micrographs of hematoxylin and eosin staining (H&E) of lung sections from mock or SARS-CoV-2 infected mice on day 3 and 10 post infection (p.i.). The green arrow points to a cluster of immune infiltrates. The red arrow indicates a cluster of brownish cells of hemosiderosis. Magnification 400 x. **c**) Iron-staining (blue) of lung sections from mock or SARS-CoV-2 infected mice on days 3 and 10p.i. Black arrows point to iron laden cells. Mock: mock infected. N=2 (mock), 4 (Day 3), 7 (Day 10), 3 (Day 35) per genotype. **e, f**) Sex-and-age matched mice were administered 350 CCID_50_ (cell culture infectious dose 50% assay) influenza A PR/8/34 H1N1 intranasally. **e**) The percentage of the body mass relative to day 0 (weighed immediately before infection). Data point: mean ± s.e.m. N= 5-6. **f**) The survival curve. N=6 per genotype. P=0.02 (Log-Rank test).

Severe COVID-19 pathogenesis is a combination of a direct cytopathic effect of SARS-CoV-2 replication and hyper-inflammation in the lung ^16^. In particular, COVID-19 fatality is strongly associated with elevated inflammatory mediators such interleukin 6 (IL-6) and tumor necrosis factor (TNF-α) etc. ^16^. We first examined viral loads and immune gene expression. The viral loads in lungs trended higher in *Ubxn3b*^−/−^ when compared to *Ubxn3b*^+/+^ mice, though they varied significantly among individuals at day 3 p.i. By day 10 p.i. the virus was almost cleared from the lung in both mouse genotypes (Suppl **Fig.s1a**). The serum cytokines IL-6, TNF-α, IL-10 and granulocyte-macrophage colony-stimulating factor (GM-CSF) were higher in *Ubxn3b*^−/−^ than those in *Ubxn3b*^+/+^ on day 3 p.i. (Suppl **Fig.s1b**), which is consistent with clinical observations in severe COVID-19 patients. The concentrations of serum interferon alpha (IFN-α), C-X-C motif chemokine ligand 10 (CXCL10) and IFN-γ were modestly upregulated, but equally between *Ubxn3b*^−/−^ and *Ubxn3b*^+/+^ mice after SARS-CoV-2 infection, suggesting a normal type I/II IFN response in *Ubxn3b*^−/−^ (Suppl **Fig.s1b**).

### UBXN3B maintains immune homeostasis during SARS-CoV-2 infection

COVID-19 fatality is strongly associated with an imbalance in immune cell compartmentalization, characterized by neutrophilia and lymphopenia ^16^. We thus analyzed neutrophils and T cells in the lung by flow cytometry and found that the total CD45^+^ immune cell counts per lung were ~10-fold higher in SARS-CoV-2-infected animals than mock-treated animals. However, there was no significant difference between *Ubxn3b*^−/−^ and *Ubxn3b*^+/+^ littermates (**Fig.2a**). Upon close examination we detected ~3 –fold increase of CD11b^+^ cells, while a modest reduction of total and CD4^+^ T cells in *Ubxn3b*^−/−^ compared to *Ubxn3b*^+/+^ mice (**Fig.2b**). Importantly, the ratio of neutrophil-to-T/B lymphocytes (N/L) in the lung was significantly higher (3.2-fold) in *Ubxn3b*^−/−^ (**Fig.2c**), which is consistent with the clinical observations in severe COVID-19 patients ^20^. This prompted us to examine more immune cell compartments in the lung and peripheral blood, and the longer impact of SARS-CoV-2 infection on immune cells. We noted ~3-fold increase in neutrophil and 10-fold decrease in B cell frequencies (**Fig.2d**); the N/L ratio was also much higher in the blood of *Ubxn3b*^−/−^ than *Ubxn3b*^+/+^ mice at day 3 p.i. (**Fig.2e**). By day 35 p.i., the total immune cells and T cells were lower, while neutrophils, macrophages/monocytes and N/L ratios were higher, in *Ubxn3b*^−/−^ than *Ubxn3b*^+/+^ lungs (**Fig.2f, g,** Suppl **Fig.s2**). Remarkably, the B cell frequencies and counts were dramatically decreased (5-20-fold) in both the lung and blood of *Ubxn3b*^−/−^ mice (**Fig.2d, f**). On day 35 p.i., *Ubxn3b*^−/−^ mice had ~40% fewer total CD45^+^ cells in the blood (**Fig.2h**) and presented splenic atrophy characterized by reduced cell density in white pulps and increased myeloid clumps in red pulps, when compared to *Ubxn3b*^+/+^ animals (Suppl **Fig.s3**). Because of the dramatic defect in B cell compartment that might lead to a weak antibody response, we thus measured the concentrations of serum anti-SARS-CoV-2 Spike and IVA nucleoprotein IgG by enzyme-linked immunosorbent assay (ELISA). Indeed, the IgG concentrations in *Ubxn3b*^−/−^ were lower than those in *Ubxn3b*^+/+^ mice (**Fig.2i**).

**Fig.2.**
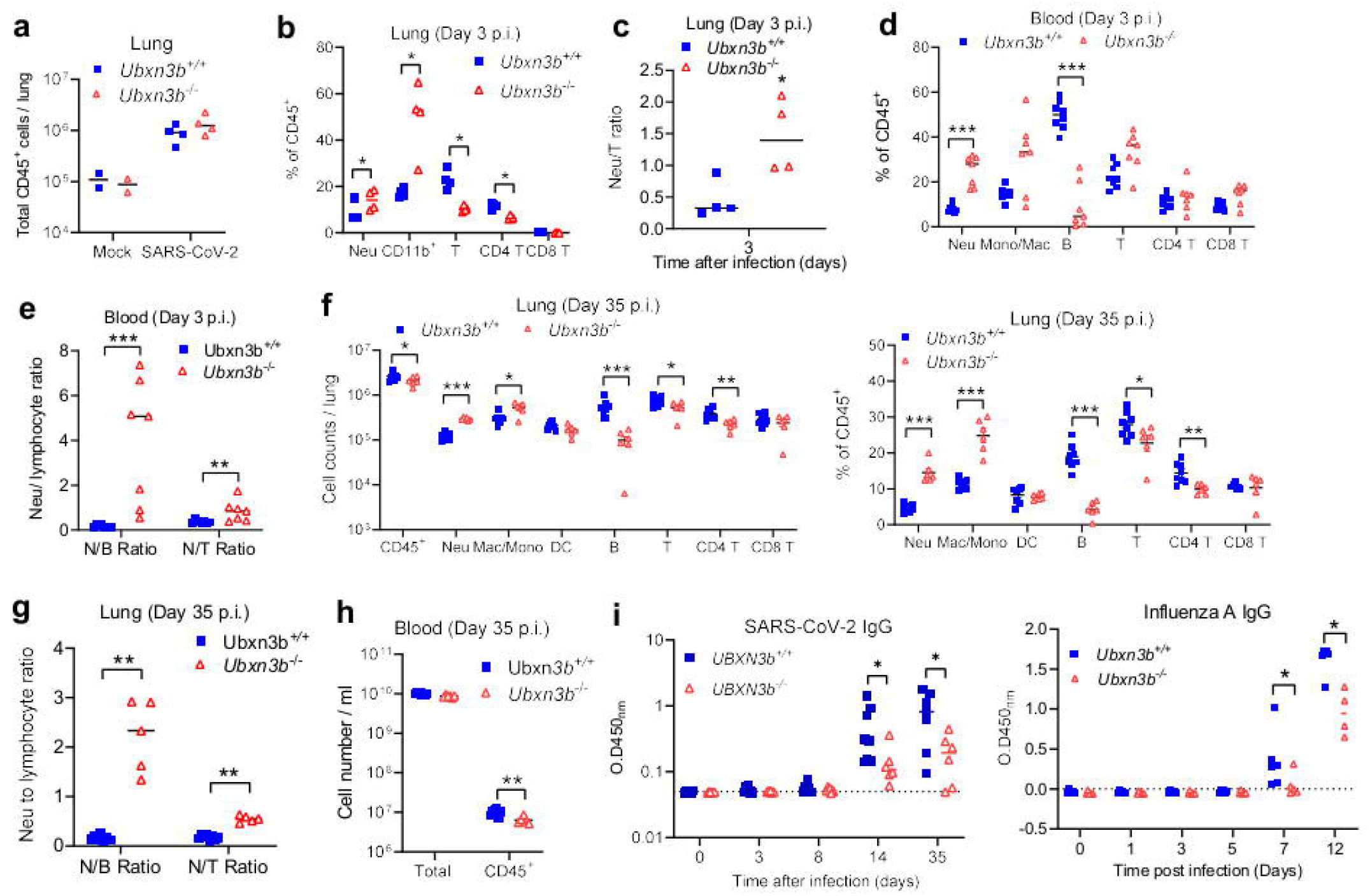
UBXN3B is essential for immune cell homeostasis during SARS-CoV-2 infection. Sex-and-age matched mice were administered 2×10^5^ plaque forming units (PFU) of SARS-CoV-2 intranasally. **a**) Total CD45^+^ cells, **b**) the percentage (relative to CD45^+^ cells) of various immune cell populations quantified by flow cytometry, **c**) the neutrophil-to-T cell ratio, in one lung of SARS-CoV-2 infected mice at day 3 post infection (p.i.). **d**) The percentage (relative to CD45^+^ cells) of various immune cell compartments, and **e**) the neutrophil-to-B/T cell ratios (N/B, N/T), in the blood at day 3 post infection (p.i.). The cell counts and percentage of various immune cell populations **f**) and **g**) the neutrophil-to-B/T cell ratios in one lung, **h**) cell counts in the blood at day 35 p.i. **i**) The concentrations of serum IgG against SARS-CoV-2 Spike and influenza A PR/8/34 H1N1 NP were quantitated by ELISA and presented as optical density at 450nm (O.D_450nm_). Neu: neutrophil, Mac/Mono: macrophage/monocyte, DC: dendritic cell. Each symbol=one mouse. *, p<0.05; **, p<0.01; ***, p<0.001 (non-parametric Mann-Whitney test). The horizontal line indicates the median of the result.

### UBXN3B maintains steady-state hematopoietic homeostasis

The above-mentioned results suggest an essential role of UBXN3B in maintenance of immune cell homeostasis during viral infection. Therefore, to test the hypothesis that dysregulated immune homeostasis is caused by *Ubxn3b*^−/−^ deficiency, we reasoned that in the steady state *Ubxn3b*^−/−^ mice should have alterations that cannot be explained by infection. Indeed, by flow cytometry, we observed a significant increase in the frequencies of myeloid cells (neutrophils, monocytes, macrophages), in the blood and spleen of *Ubxn3b*^−/−^ compared to *Ubxn3b*^+/+^ mice. Although the T cell frequencies were slightly lower or not different, its counts were significantly lower in both tissues of *Ubxn3b*^−/−^ than *Ubxn3b*^+/+^ mice (**Fig.3a, b**). The total CD45^+^ count per spleen was ~3-fold lower in *Ubxn3b*^−/−^ (**Fig.3b**). Of note, *Ubxn3b*^−/−^ mice had over 10 times lower B cell counts and frequencies in both tissues of knockout mice (**Fig.3a, b**). Although the magnitudes of difference in T cell frequency varied with tissues, the N/T ratios were uniformly much higher in all tissues of *Ubxn3b*^−/−^ when compared to *Ubxn3b*^+/+^ mice (**Fig.3c**). Next, we asked if STING has a role in steady state hematopoiesis because our recent study demonstrated a role of UBXN3B in activating STING signaling ^15^. The results show that the blood immune cell compositions were similar between wild type and *Sting*^−/−^ mice (Suppl **Fig.s4**), suggesting a STING-independent role of UBXN3B in hematopoiesis.

**Fig.3.**
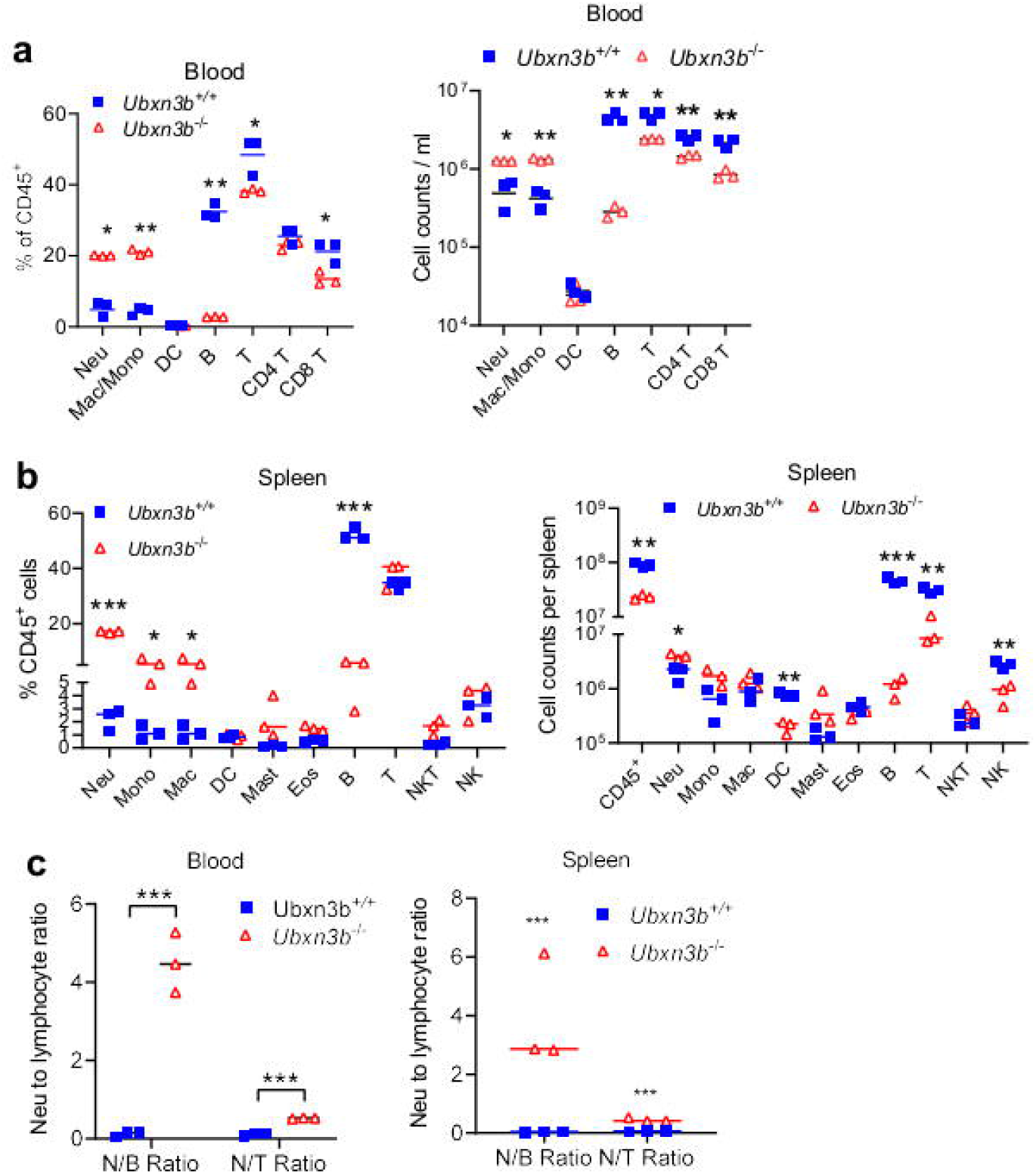
UBXN3B is essential for steady-state immune cell homeostasis. The percentage (relative to CD45^+^ cells) of various immune cell populations and cell counts were quantified by flow cytometry in the **a**) blood and **b**) spleen of specific pathogen-free littermates. **c**) The neutrophil to B/T cell ratios. Neu: neutrophil, Mac/Mono: macrophage/monocyte, DC: dendritic cell, Mast: mast cell, Eos: eosinophil, NK: natural killer, NKT: natural killer T cells. Each symbol=one mouse. The horizontal line indicates the median of the result. *, p<0.05; **, p<0.01; ***, p<0.001 (two-tailed Student’s *t*-test).

Hematopoiesis involves a global change of gene expression controlled by cell-intrinsic transcription factors and epigenetic modifiers, and cell-extrinsic factors such as cytokines, chemokines, growth factors, and interactions with osteoblasts, endothelial cells, reticular cells and stromal cells in its bone marrow niche ^3^. To investigate if the B cell defect in *Ubxn3b*^−/−^ is cell-intrinsic or extrinsic, we performed reciprocal bone marrow transplantation. Firstly, we transferred Cre^+^ *Ubxn3b* ^flox/flox^ bone marrow (CD45.2) to irradiated wild type (WT, CD45.1) recipient mice. Thirty days after transplantation, the recipient WT mice were treated with either corn oil (designated *Ubxn3b*^+/+^BM–WT) or tamoxifen (dissolved in corn oil) to induce *Ubxn3b* deletion in the bone marrow (BM) (designated *Ubxn3b*^−/−^ BM–WT). We confirmed that > 99% of blood B cells /neutrophils/monocytes, and >82% of T cells were derived from the CD45.2 donor at 45 days after transplantation (Suppl **Fig.s5a**), indicating successful irradiation and immune reconstitution. We noted an unusually high neutrophil ratio in *Ubxn3b*^+/+^ BM–WT mice at day 15 than regular *Ubxn3b*^+/+^ mice (~32% versus ~10%), and it was back to normal by day 30 (**Fig.4a,b,** Suppl **Fig.s5b**). This is likely because of faster reconstitution of neutrophils than B cells after BMT. Nonetheless, the B cell cellularity in the chimeric *Ubxn3b*^−/−^ BM–WT mice was consistently much lower than that in *Ubxn3b*^+/+^ BM–WT mice throughout the study period (**Fig.4a-c**). Moreover, *Ubxn3b*^−/−^ BM–WT mice were more vulnerable to SARS-CoV-2 infection, typified by more iron-laden cells than *Ubxn3b*^+/+^ BM–WT mice were in the lung at day 7 p.i (**Fig.4d**). Conversely, *Ubxn3b* ^flox/flox^ or Cre^+^ *Ubxn3b* ^flox/flox^ mice (CD45.2) were irradiated and transplanted with WT (CD45.1) bone marrow. Thirty days after transplantation, the recipient mice were treated with tamoxifen, resulting in chimeric WT *BM-Ubxn3b*^+/+^ and WT BM–*Ubxn3b*^−/−^ mice. The B cell numbers were comparable between the two groups (Suppl **Fig.s6**). These data suggest that UBXN3B plays a cell-intrinsic role in controlling B cell development and hematopoietic UBXN3B is critical for restricting SARS-CoV-2 pathogenesis.

**Fig.4.**
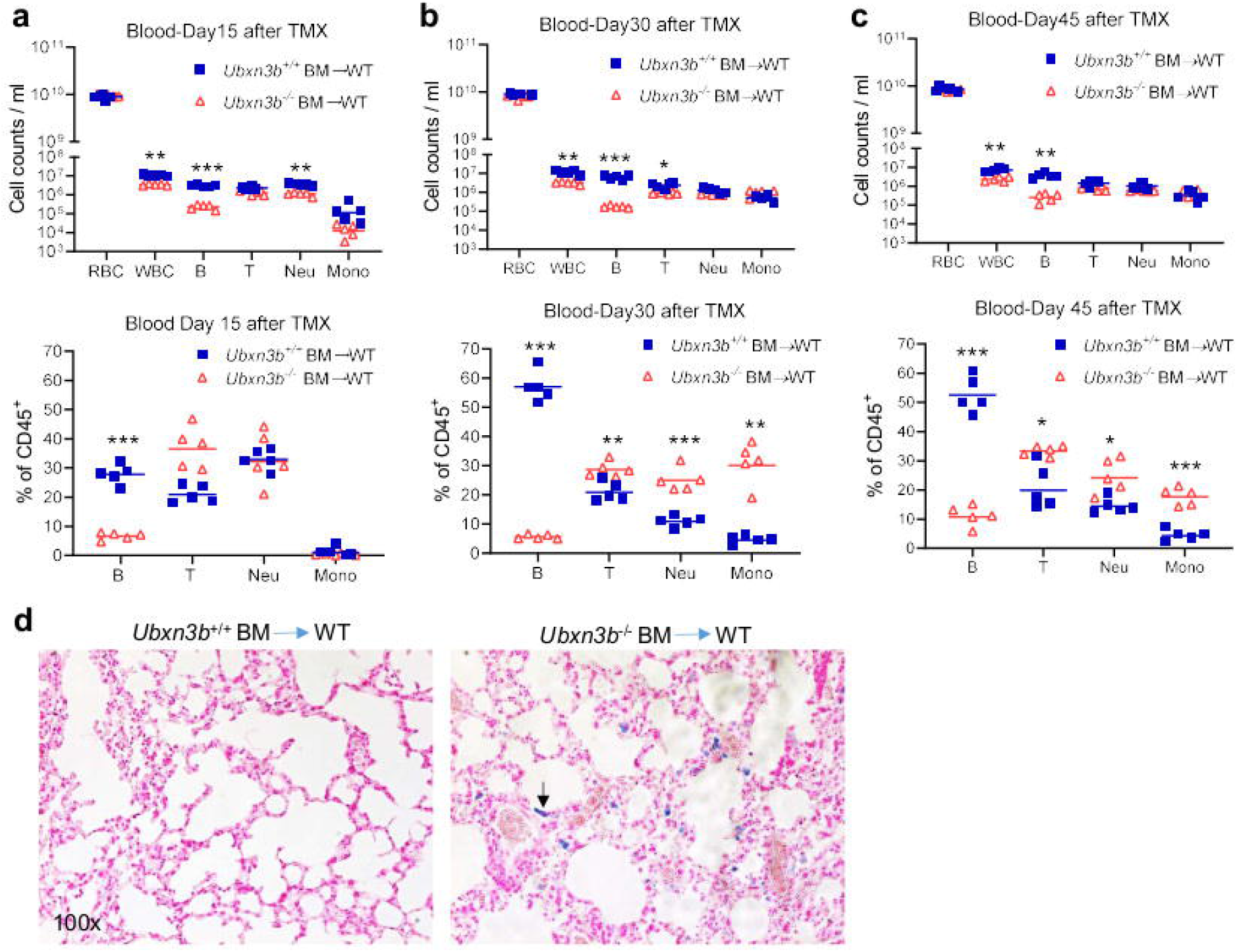
The essential role of UBXN3B for B cell development is cell-intrinsic. Irradiated wild type (WT, CD45.1) recipient mice were transplanted with *Cre*+ *Ubxn3b*^f/f^ bone marrow (CD45.2). The mice were then treated with tamoxifen (TMX) to delete Ubxn3b in hematopoietic cells (designated *Ubxn3b*^−/−^ BM–WT) or corn oil (designated *Ubxn3b*^+/+^ BM–WT). The cell counts and percentage (relative to CD45^+^ cells) of various immune cell populations in the blood were quantified by flow cytometry at days **a**) 15, **b**) 30, **c**) 45 after completion of the TMX treatment. **d**) Iron-staining (blue) of lung sections at day 7 p.i. The black arrow points to iron-laden cells. Magnification: 100x. N=5. Each symbol=one mouse. The horizontal line indicates the median of the result. *, p<0.05; **, p<0.01; ***, p<0.001 (two-tailed Student’s *t*-test).

### UBXN3B controls B lymphopoiesis by maintaining BLNK protein stability and pre-BCR signaling

The aforementioned results suggest that UBXN3B likely regulates B cell development. To this end, we quantitated terminally differentiated immune cells in the bone marrow. Among all live cells (after lysis of red blood cells), neutrophil was the most abundant, then B cell. The percentage of B cells was ~6-fold lower, while the frequency of neutrophils was moderately higher, in *Ubxn3b*^−/−^ than *Ubxn3b*^+/+^ bone marrow (**Fig.5a,** Suppl **Fig.s7a**). These results suggest that dysregulated hematopoiesis in *Ubxn3b*^−/−^ is due to a defect in B lymphopoiesis, which we tested by assessing all the stages of B cell development. Of note, the percentage of large and small precursor BII (pre-B), immature B (imm-B) and mature B (recirculating B) fractions was significantly lower in *Ubxn3b*^−/−^ than *Ubxn3b*^+/+^, while that of progenitor B (pro-B) and precursor BI (pre-BI) fractions was the same (**Fig.5b,** Suppl **Fig.s7b**), suggesting that UBXN3B is essential for pre-BI transition to pre-BII, the first checkpoint. Next, we examined stem cells and other lineage progenitors. Total HSCs (Lin^−^ Sca^+^ Kit^+^) contains two populations, long-term HSCs, which are capable of self-renewal but are quiescent at steady state, and short/mid-term multipotent HSCs (also known as MPPs), which are capable of differentiating into lineage-committed progenitors. The LSK and MPP percentage was modestly decreased in *Ubxn3b*^−/−^ when compared to *Ubxn3b^+/+^*. The frequency of lineage-committed common lymphoid progenitors (CLPs) was also reduced, while the common myeloid progenitors (CMPs) trended higher in *Ubxn3b*^−/−^ (**Fig.5c,** Suppl **Fig.s8**).

**Fig.5.**
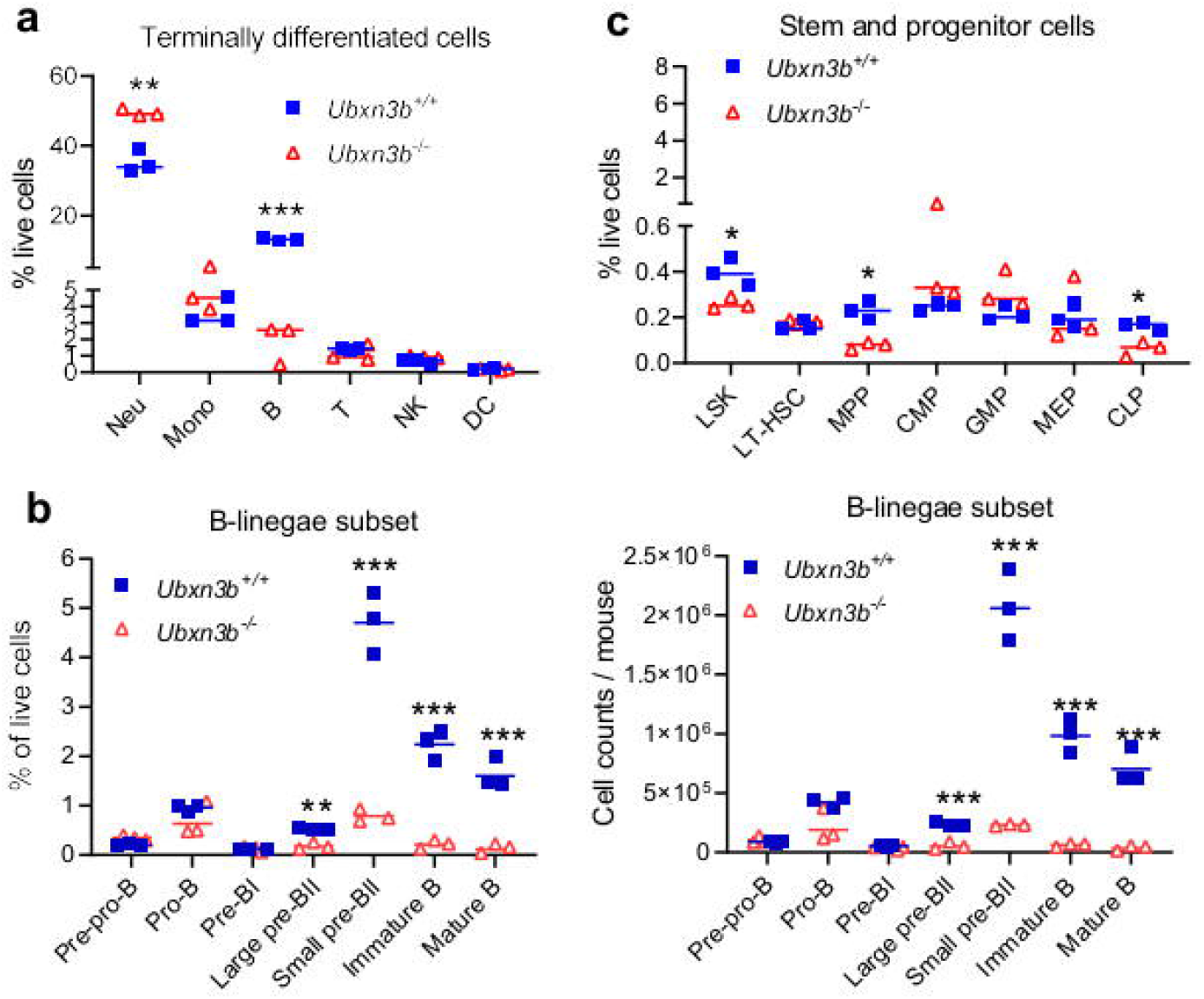
UBXN3B is essential for pre-BI transition to pre-BII. The frequencies of **a**) terminally differentiated immune cells and **b**) stem cells/progenitors, quantified by flow cytometry (relative to live cells after lysis of red blood cells). **c**) The frequencies and cellularity of B lineage subsets in the bone marrow of specific pathogen-free littermates. Neu: neutrophil, Mono: monocyte, DC: dendritic cell, NK: natural killer, LSK: Lin^−^ Sca^+^ Kit^+^, LT-HSC: long-term hematopoietic stem cell, ST-HSC: short-term multipotent HSC (also known as MPP), CMP: common myeloid progenitor, CLP: common lymphoid progenitor, GMP: granulocyte–macrophage progenitor, MEP: megakaryocyte–erythroid progenitor, pre-pro-B: pre-progenitor B, pro-B: progenitor B, pre-B: precursor B. Each symbol=one mouse. The horizontal line indicates the median of the result. *, p<0.05; **, p<0.01; ***, p<0.001 (two-tailed Student’s *t*-test).

The abovementioned results demonstrate that UBXN3B is essential specifically for early B cell development, and this is cell-intrinsic. Therefore, we sorted early bone marrow B fractions and quantified by qRT-PCR the mRNA expression of well-established transcription factors for hematopoiesis, several of which are B lineage-specific/dominant transcription factors, including early B cell factor (EBF1), paired box protein 5 (PAX5) myocyte enhancer factor (MEF2C), and Ikaros family zinc finger 1/3 (IKZF1/3) ^21^. *Ebf1*, *Pax5* and *Ikzf3* (encodes Aiolos) mRNA levels were dramatically induced (>100 fold) in pro-B, pre-B and mature B, when compared to those in pre-pro-B cells. Of note, *Ikzf3* was decreased by 5-fold and *Ikzf1* (encodes Ikaros) was modestly reduced but only transiently in *Ubxn3b*^−/−^ pro-B cells, when compared to that in *Ubxn3b*^+/+^ pro-B cells (Suppl **Fig.s9a**). We also checked B cell surface marker genes (*Cd19*, *Cd79a* and *Il7ra*) and observed no significant difference (Suppl **Fig.s9b**). These data suggest that the transcription factor expression in general is intact in *Ubxn3b*^−/−^ B lineage.

Because our data have shown that UBXN3B is essential for the transition from pre-BI to large pre-BII (checkpoint 1), we postulated that UBXN3B might regulate pre-B cell receptor (BCR) signaling. Pre-BCR signaling plays several important roles, including allelic exclusion, negative/positive selection, and proliferation of large pre-BII ^22^. A Pre-BCR comprises an Ig μ heavy chain (μH) and a surrogate light chain (SLC); the latter is transiently robustly induced in pro-B and pre-BI, but rapidly down-regulated in large pre-BII to allow for BCR recombination and expression ^23^. Indeed, *Vpreb* expression (V-set pre-B cell surrogate light chain) was up-regulated by >150 times in pro-B when compared to pre-pro-B, further in pre-BI cells, then was suppressed in large pre-BII, but barely detected in recirculating B and other fully differentiated lineages. We observed a modest increase in *Vpreb* mRNA in pro-B, pre-BI and large pre-BII of *Ubxn3b*^−/−^, compared to that of *Ubxn3b*^+/+^ mice (**Fig.6a**). However, the surface Vpreb1 protein abundance was comparable between Vpreb1^+^ (pro-B, pre-BI, large pre-BII cells) *Ubxn3b*^−/−^ and *Ubxn3b*^+/+^ cells (**Fig.6b)**. Of note, *Ubxn3b* expression was higher in the B lineage than T/neutrophil/monocytes, being the highest in pre-BI, coincident with its essential role at the first checkpoint (**Fig.6a**). Next, we attempted to obtain a full picture of the pathways regulated by UBXN3B. To this end, we performed single cell RNA sequencing (scRNAseq) on all bone marrow B fractions (excluding mature, recirculating B), HSCs and progenitors (CLPs, GMPs and MEPs). We analyzed the differentially expressed genes (DEGs) in SLC-high (SLC^hi^) B subsets (**Fig.6c**) and identified 49 down-regulated genes with an average count per cell >1 and a p<0.1 (Log2 fold range −0.7 to −1.7) in *Ubxn3b*^−/−^ SLC^hi^ cells. Of note, 10 genes (~20% of total down-regulated DEGs) were related to the cell cycle/mitosis/DNA replication pathways (Suppl **Table s1, Fig.6d**). The opening of IgK gene locus and recombination by RAG1/2 downstream of pre-BCR is essential for B lymphopoiesis ^24,25^. Intriguingly, *Igkc* was downregulated too (Log2 = −1.3, p=0.00025). Fifty genes were upregulated, including three SLC genes (*Vpreb1/2* and *Igll1*) (Suppl **Table s1**), consistent with the qRT-PCR result (**Fig.6a**). In *Ubxn3b*^−/−^ SLC-low (SLC^lo^) B cells, 23 genes were significantly down-regulated (Log2 range −0.7 to −2.9), three of which were BCR genes (*Iglc1*, *Iglc2*, *Ighd*); but no significant pathway was identified (Suppl **Table s2**). We noted only 5-7 significantly down-regulated genes and no significantly enriched pathways in HSCs or GMPs (Suppl **Table s3, 4**). These data suggest that UBXN3B likely regulates pre-BCR downstream signaling components. To this end, we first examined these protein expression. B-cell linker (BLNK/SLP65) expression was gradually reduced from pre-pro-B to large pre-B cells; it was much lower in *Ubxn3b*^−/−^ than that in corresponding *Ubxn3b*^+/+^ B fractions. The level of Bruton’s tyrosine kinase (BTK), phospholipase C gamma 2 (PLC-γ2), transcription factors forkhead box protein O1 (FoxO1) and CCAAT-enhancer-binding protein α (C/EBPα) remained constant and similar between all *Ubxn3b*^+/+^ and *Ubxn3b*^−/−^ B fractions (**Fig.6e**). *Blnk* mRNA expression was similar between *Ubxn3b*^+/+^ and *Ubxn3b*^−/−^, but lower in SLC^lo^ than SLC^hi^ B cells (**Fig.6f**). Next we checked if pre-BCR signaling is defective by measuring calcium influx. We purified bone marrow B fractions including SLC^hi^ pro-B/ pre-BI/large pre-BII, SLC^lo^ small pre-BII and immature B, and mature B fractions by FACS; stimulated (pre-) BCR signaling with an anti-IgM μH antibody, and monitored calcium flux over a time course by flow cytometry. The anti-IgM μH antibody significantly increased calcium influx in all *Ubxn3b*^+/+^ B fractions, which was impaired in *Ubxn3b*^−/−^ cells (**Fig.6g**). Of note, the difference in the calcium flux and BLNK protein level between *Ubxn3b*^−/−^ and *Ubxn3b*^+/+^ mature B cells became smaller. These data suggest that UBXN3B maintains BLNK protein stability specifically during early B development.

**Fig.6.**
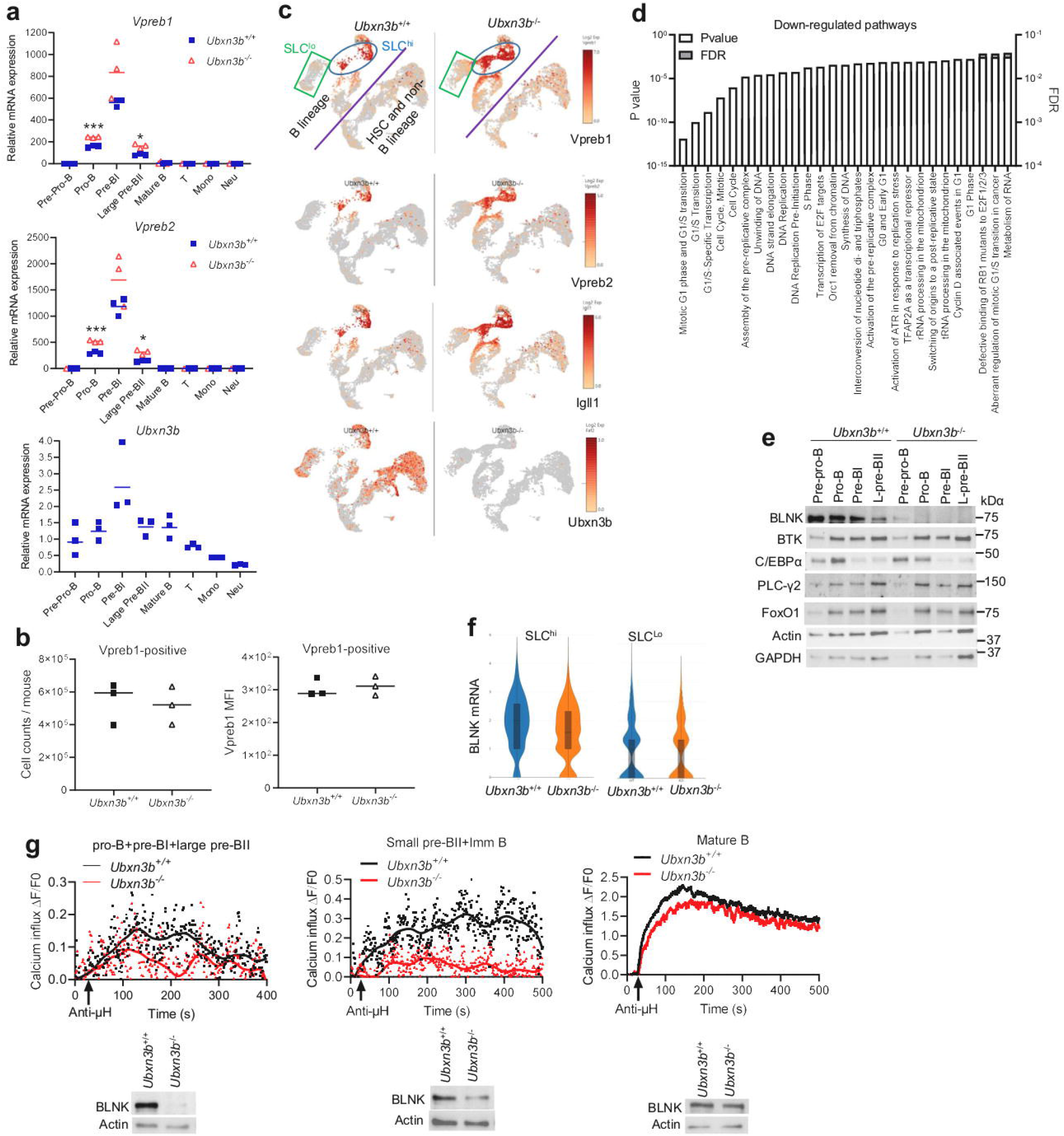
UBXN3B maintains BLNK protein level and pre-BCR signaling. **a**) qRT-PCR quantification of gene expression in bone marrow B lineage subsets of specific pathogen-free littermates. Pre-pro-B: pre-progenitor B, pro-B: progenitor B, pre-B: precursor B, Neu: neutrophil, Mono: monocyte. **b**) The cellularity of surface Vpreb1^+^ cells and mean fluorescence intensity (MFI). Each symbol=one mouse. The horizontal line indicates the median of the result. *, p<0.05; **, p<0.01; ***, p<0.001 (two-tailed Student’s t-test). **c**) The Uniform Manifold Approximation and Projection (UMAP) of surrogate light chain (SLC) gene expression by scRNA-seq. Cells in the oval express a high level of SLC (SLC^hi^), while cells in the rectangle express a low level of SLC (SLC^lo^). **d**) The most significant pathways for the down-regulated genes in *Ubxn3b*^−/−^ SLC^hi^ cells, when compared to *Ubxn3b*^+/+^ cells. FDR: false discovery rate. **e**) Immunoblots for the indicated proteins in bone marrow B fractions. L: large. **f**) The violin plots of *Blnk* transcript levels by scRNAseq in bone marrow SLC^hi^ and SLC^lo^ B cells. **g**) The ratio of ΔF (the difference of calcium load between any a given time after anti-IgM μH treatment and time point zero F0) to F0. Each dot represents the ratio of mean ΔF/F0 of all the cells recorded at a given time (every second). Below each chart is the immunoblot of BLNK. **c-g**) represent the results from three mice.

## DISCUSSION

The HSC differentiation cascade must be finely regulated to enable timely production of the right number and type of mature cells, i.e. homeostasis, disruption of which may lead to a pathological state, such as autoimmunity, immunodeficiency, cancer etc. As an example, lymphopenia and a skewing myeloid-to-lymphoid ratio in the elderly may contribute to inflammaging and impaired immunity ^26^. In this study, we have discovered a novel and essential role of UBXN3B in maintenance of hematopoietic homeostasis, of note, B cell development, and control of immunopathogenesis of respiratory viral diseases.

### The mechanism of UBXN3B action in RNA virus pathogenesis

Mechanistically, UBXN3B might control RNA virus infection by regulating STING signaling and type I IFN responses ^15 27^. However, expression of type I IFNs is normal in *Ubxn3b*^−/−^ mice, so is steady-state hematopoiesis in *Sting*-deficient mice. These results suggest that the primary function of UBXN3B during RNA virus infection is independent of STING and that dysregulated hematopoiesis may be the main contributor to failure of viral clearance and prolonged immunopathology in *Ubxn3b*^−/−^ mice. Indeed, although belonging to very different families of RNA viruses, both SARS-CoV-2 and IVA elicit immunopathology in the lung, including massive immune infiltrates and elevated levels of systemic pro-inflammatory mediators ^17^. In particular, COVID-19 fatality is strongly associated with neutrophilia and lymphopenia ^16^, which is partly recapitulated in *Ubxn3b*^−/−^ mice. Intriguingly, regardless of viral loads, all *Ubxn3b*^−/−^ mice present a similar degree of hemosiderosis at days 3 and 10 p.i., while none of *Ubxn3b*^+/+^ mice have evident hemosiderosis, suggesting that the severe lung damage in *Ubxn3b*^−/−^ mice is primarily caused by immunopathology. On the other hand, heightened immunopathology and tissue damage persists in *Ubxn3b*^−/−^ mice even after viral clearance (day 10 post SARS-CoV-2 inoculation), suggesting that immunopathogenesis is disassociated from viral replication. Consistent with this notion, the hyperinflammatory phase of clinical COVID-19 generally happens after the viral load peak, with a few infectious viral particles ^28^. At the post clearance stage, a high N/L ratio may sustain inflammation, and thus is correlated with poor prognosis of severe COVID-19 patients ^29 20^. The N/L ratio is also the most reliable biomarkers of chronic inflammatory conditions, such as type II diabetes ^30^, cardiovascular disease ^31^, and aging ^32^ etc. These are actually significant risk factors for COVID-19 mortality^16^. These conditions are characterized by a low-grade pro-inflammatory and an “immunosenescence”-like immune state that is unable to clear viruses ^20^. In this regard, UBXN3B deficiency might resemble the aging immune state.

### The mechanism of UBXN3B action in B cell development

Although our initial pursuit with UBXN3B focused on RNA virus pathogenesis, the unexpected and significant phenotype in B cell compartment prompted us to delve into B lymphopoiesis. Mechanistically, this defect is cell intrinsic because reconstitution of the hematopoietic system of *Ubxn3b*^−/−^ mice with WT bone marrow restores B cellularity, while reverse transplantation fails to do so. However, expression of cell-intrinsic B-lineage transcription factors (Pax5, Ebf1, Myb, Ikzf1) is largely normal except for a transient downregulation of Ikzf1/3 in pro-B only, which seems unlikely accountable for a significant defect in the B cell compartment in *Ubxn3b*^−/−^ mice. Of note, the cellularity of pre-BI and B progenitors remains normal, until the large pre-BII stage in *Ubxn3b*^−/−^ mice, suggesting a failure of transition from pre-BI to large pre-BII, also known as the first checkpoint. This stage requires a transient yet essential pre-BCR signaling to drive allelic exclusion, negative/positive selection, and proliferation of large pre-BII ^22^. Indeed, in *Ubxn3b*^−/−^ SLC^hi^ B subset (primarily pre-BI) ^33^, the down-regulated genes are enriched in the cell cycle/mitosis/DNA replication pathways. However, in *Ubxn3b*^−/−^ SLC^Lo^ B fraction (predominantly small pre-BII and immature B) ^33^, there is no significantly down-regulated pathway. These data suggest that UBXN3B is associated with a developmental stage-specific signaling pathway, e.g., pre-BCR. Indeed, our results demonstrate that UBXN3B is essential for maintenance of BLNK protein stability during the early stage of B development. BLNK is a scaffold protein that is essential for assembly of a macromolecular complex comprising BTK, PLC-γ etc. of the (pre-) BCR pathway ^34^. BLNK acts at B220^+^ CD43^+^ pro-B transition to B220^+^ CD43^−^ pre-B ^35,36^. Similarly, with more surface markers, we show that UBXN3B is essential for B220^+^ CD43^+^ pre-BI transition to B220^+^ CD43^+^ large pre-BII and then proliferation of B220^+^ CD43^−^ small pre-BII. However, UBXN3B is no longer essential for BLNK protein stability and BCR signaling in mature B cells. This is likely because that a few *Ubxn3b*^−/−^ cells at the early stage still express sufficient BLNK and finally differentiate into mature B. Considering that UBXN3B is also expressed by mature B cells, it is plausible that UBXN3B may dependent on an early B stage-specific cellular factor to maintain a high BLNK level.How does UBXN3B regulate BLNK protein level? Many UBXNs including UBXN3B are known to interact with p97 and multiple E3 ligases, thus controlling newly synthesized protein quality, and regulating protein turnover^8,9^. Thus, by interfacing different E3 ligases under different physiological, developmental or tissue contexts, UBXNs could participate in multiple cellular functions. Indeed, UBXN3B works with tripartite motif-containing 56 (TRIM56) to ubiquitinate and activate STING during DNA virus infection ^15^. In early B development, UBXN3B could inhibit a specific E3 ligase that mediates BLNK turnover. In support of this concept, we recently showed that UBXN6 inhibits degradation of phosphorylated tyrosine kinase 2 (TYK2) and type I/III interferon receptor activated by type I/III IFNs^37^.

In summary, our results demonstrate that UBXN3B is essential for maintenance of hematopoietic homeostasis and in particular B lymphopoiesis during steady state and viral infection. Aberrant immune compartmentalization associated with UBXN3B deficiency may predispose an individual to persistently heightened immunopathology during viral infection. Future work will address how UBXN3B regulates BLNK stability.

## MATERIALS AND METHODS

### Mouse models

The mouse line with the exon 1 of Ubxn3b flanked by two LoxP sites (Ubxn3b^flox/flox^) were generated via homologous recombination by Dr. Fujimoto at Nagoya University ^38^. The homozygous Ubxn3b^flox/flox^ were then crossed with homozygous tamoxifen-inducible Cre recombinase-estrogen receptor T2 mice (The Jackson Laboratory, Stock # 008463) to generate Cre^+^ *Ubxn3b*^flox/flox^ littermates. To induce Ubxn3b deletion, > 6-weeks old mice were injected with 100□μl of tamoxifen (10□mg /□ml in corn oil) (Sigma, #T5648) via intraperitoneal (i.p.) every 2 days for a total duration of 8 days (4 doses). Successful deletion of Ubxn3b was confirmed in our recent study ^12^. A half of Cre^+^ *Ubxn3b*^flox/flox^ litters were treated with tamoxifen and designated *Ubxn3b*^−/−^; the other half were treated with corn oil only and designated *Ubxn3b*^+/+^. Mice were allowed to purge tamoxifen for at least 4 weeks before any infection or analyses was performed. B6.SJL-Ptprc^a^ Pepc^b^/BoyJ (Stock No. # 002014) is a congenic strain used widely in transplant studies because it carries the differential pan leukocyte marker Ptprc^a^, commonly known as Cd45.1 or Ly5.1. All experiments were performed in accordance with relevant guidelines and regulations approved by the Institutional Animal Care and Use Committee at the University of Connecticut and Yale University.

### Antibodies, Cell lines and Viruses

A rabbit anti-BLNK mAb (Clone D3P2H, Cat #36438), anti-ß-Actin mAb (Clone D6A8, Cat # 8457), anti-phospho-BLNK mAb (Thr152) (Clone E4P2P, Cat #62144), anti-GAPDH (Clone D16H11, Cat # 5174), anti-BTK mAb (Clone D3H5, Cat # 8547), anti-phospho-BTK mAb (Tyr223) (Clone D1D2Z, Cat # 87457), anti-Syk mAb (Clone D3Z1E, Cat # 13198), anti-phospho-Zap-70 (Tyr319)/Syk (Tyr352) mAb (Clone 65E4, Cat # 2717), anti-MEK1/2 mAb (Clone D1A5, Cat # 8727), and anti-phospho-MEK1/2 mAb (Ser221) (Clone 166F8, Cat # 2338) were purchased from Cell Signaling Technology (Danvers, MA 01923, USA). Human embryonic kidney 293 cells transformed with T antigen of SV40 (HEK293T, # CRL-3216) and Vero cells (monkey kidney epithelial cells, # CCL-81) were purchased from American Type Culture Collection (ATCC) (Manassas, VA20110, USA). These cell lines are not listed in the database of commonly misidentified cell lines maintained by ICLAC. Cells were grown in DMEM supplemented with 10% fetal bovine serum (FBS) and antibiotics/antimycotics (Life Technologies, Grand Island, NY 14072 USA).We routinely added MycoZAP (Lonza Group, Basel, Switzerland) to cell cultures prevent mycoplasma contamination.

SARS-CoV-2 (NR-52281, Isolate USA-WA1/2020) was provided by BEI Resources (funded by National Institute of Allergy and Infectious Diseases and managed by ATCC, United States). The full-length human ACE2 [Accession No: NM_021804.2] cDNA was inserted into pAV-EGFP-CMV/FLAG and Ad5 viruses were prepared by Vector Builder Inc. (Chicago, IL 60609, USA).

### Concentration of SARS-CoV-2

The virus was grown in Vero cells for 72hrs, and the culture medium was cleared by brief centrifugation. A PEG-it Virus Precipitation Solution (Cat# LV810A-1, System Biosciences, Palo Alto, CA 94303, USA) was added to 40ml of virus culture at a 1:4 ratio, incubated overnight at 4°C. The mixture was centrifuged at 1500xg for 30 min, and the resulting pellet was suspended in 1ml of DMEM medium. In parallel, Vero cell culture medium without virus was processed in the same way and used for mock infection.

### Plaque-Forming Assay

Quantification of infectious viral particles in sera or homogenized tissues was performed on Vero cell monolayer ^39^. Briefly, viral samples were incubated with confluent Vero cells (6-well plate) at 37□°C for 2□hr. The inoculum was then removed and replaced with 2□ml of DMEM complete medium with 1% SeaPlaque agarose (Cat. # 50100, Lonza). The cells were incubated at 37 °C, 5% CO_2_ for 3 days, and on the fourth day the cells were stained with Neutral Red (Sigma-Aldrich) overnight.

### Mouse Infection and Monitoring

Mice were administered intranasally 2×10^8^ plaque forming units (PFU) of Ad5-hACE2, after 5 days then intranasally inoculated with 2×10^5^ PFU of SARS-CoV-2 or mock. Three hundred and fifty CCID_50_ (cell culture infectious dose 50% assay) of Influenza A PR8/34 H1N1 strain was administered to mice by intranasal instillation in 40μl of sterile phosphate buffered saline. The body mass of individual mice was weighed on the day of infection (Day 0) as a baseline. The percentage change in an animal was calculated as 100 × (Day n-Day 0)/Day 0, where n defines the length of infection (in days).

### Bone Marrow Transplantation

Eight weeks or older wild type (WT, B6, Cd45.1, recipient) male mice were irradiated at a lethal dose (900 rad) with a Gammacell-40 irradiator once, and transplanted with ERT2-Cre^+^-Ubxn3b^flox/flox^ bone marrow (BM) cells (donor, Cd4.2) intravenously. Thirty days after transplantation, a half of the mice were administered 100μl of tamoxifen (10□mg /□ml in corn oil) (Sigma, #T5648) via intraperitoneal injection (i.p.) every 2 days for a total duration of 8 days (4 doses) (designated *Ubxn3b*^−/−^ BM–WT). The other half was treated with corn oil in the same way (designated *Ubxn3b*^+/+^ BM–WT). Conversely, *Ubxn3b* ^flox/flox^ or ERT2-Cre^+^ *Ubxn3b* ^flox/flox^ mice were irradiated and transplanted with WT BM. Thirty days after transplantation, all the recipient mice were treated with tamoxifen, resulting in chimeric WT *BM–Ubxn3b*^+/+^ and WT *BM-Ubxn3b*^−/−^ mice. Fifteen to forty five days after the last dose of tamoxifen, immune cells were analyzed by flow cytometry and/or mice were infected with SARS-CoV-2.

### Tissue Histology

Tissues were fixed in 4% paraformaldehyde (PFA), embedded in paraffin, cut into 4μM-thick sections, immobilized to glass slides, decalcified, and processed for hematoxylin and eosin staining. Arbitrary arthritic disease scores (on a 1–5 scale with 1 being the slightest, 5 the worst) were assessed using a combination of histological parameters, including exudation of fibrin and inflammatory cells into the joints, alteration in the thickness of tendons or ligament sheaths, and hypertrophy and hyperlexia of the synovium^40^ in a double-blinded manner.

Hemosiderosis was evaluated by iron staining (Prussian Blue stain) (Cat. # ab150674, from Abcam, Cambridge, CB2 0AX, UK). Lungs were fixed in 4% PFA, embedded in paraffin, cut into 4μM-thick sections, immobilized to glass slides, deparaffinized in xylene, rinsed with 100% ethanol, hydrated progressively in 95%, 70% ethanol and distilled water. The slides were incubated in Iron Stain Solution (1:1 of potassium ferrocyanide solution to hydrochloric acid solution) for 3 min at ambient temperature, rinsed thoroughly in distilled water, stained in Nuclear Fast Red Solution for 5 minutes, rinsed again with distilled water 4 times, dehydrated in 95% alcohol followed by absolute alcohol, and finally mounted in synthetic resin. The slides were assessed with an Accu-Scope microscope EXI-310 and images were acquired by an Infinity II camera and software.

### Flow cytometry and Fluorescence-Activated Cell Sorting (FACS)

Flow and FACS was performed according to our published study ^41^. Mouse tissues were minced with a fine scissor and digested in 4 mL of digestion medium [20 mg/mL collagenase IV (Sigma-Aldrich, St. Louis, MO, USA), 5 U/mL dispase (StemCell, Cambridge, MA, USA), and 50 mg/mL DNase I mix (Qiagen, Germantown, MD, USA) in complete RPMI1640 medium] at 37 °C for 4 hrs. The lysate was filtrated with a 40μm cell strainer. Cells were then pelleted down by centrifugation at 500 × g for 5 min. The red blood cells in the cell pellet were lysed three times with a lysis buffer (Cat. # 420301 from BioLegend, San Diego, CA 92121, USA). Cells were suspended in FACS buffer and stained for 30 min at 4 °C with the desired antibody cocktails (BioLegend, San Diego, CA, US) of APC-Fire 750-anti CD11b (Cat. # 101261, clone M1/70), Alexa Fluor 700-anti Ly-6G (Cat. # 127621, clone 1A8), Brilliant Violet 421-anti CD11c (Cat. # 117343, clone N418), PerCP-Cy5.5-anti MHC II (Cat. # 107625, clone M5/114.15.2), PE-anti Tetherin (PCDA1) (Cat. # 12703, clone 10C2), Brilliant Violet 510-anti F4/80 (Cat. # 123135, clone BM8), APC-anti CD68 (Cat. # 137007, clone FA-11), PE-Dazzle 594-anti CD3 epsilon (Cat. # 100347, clone 145-2C11), Brilliant Violet 711-anti CD4 (Cat. # 100557, clone RM4-5), Brilliant Violet 570-anti CD8a (Cat. # 100739, clone 53-6.7), Brilliant Violet 650 anti-CD161 (NK1.1) (Cat. # 108735, clone PK136), FITC anti-CD117 (cKit) (Cat. # 105805, clone 2B8), PE anti-erythroid cells (Cat. # 116207, clone TER-119), Brilliant Violet 711-anti CD115 (Cat. # 135515, clone AFS98), FITC-anti CD25 (Cat. # 102005, clone PC61), Zombie UV (Cat. # 423107), PE-Cy7-anti CD45 (Cat. # 103113, clone 30-F11), TruStain FcX-anti CD16/32 (Cat. # 101319, clone 93), APC anti-CD127 (IL-7Ra) (Cat. # 135011, clone A7R34), PE-Dazzle 594 anti-Sca-1 (Ly-6A/6E) (Cat. # 108137, clone D7), PE-Cy5 anti-Flt-3 (CD135) (Cat. # 135311, clone A2F10), Brilliant Violet 421-anti CD34 (Cat. # 119321, clone MEC14.7), PE anti-CD16/32 (Cat. # 101307, clone 93), Brilliant Violet 711-anti IgM (Cat. # 406539, clone RMM-1), Brilliant Violet 421-anti CD45R (B220) (Cat. # 103239, clone RA3-6B2), Alexa Fluor 700-anti CD19 (Cat. # 115527, clone 6D5), PE-Cy7 anti-CD93 (Cat. # 136505, clone AA4.1) and Lin- (anti-CD4, CD8, CD11b, CD11c, Gr1, NK1.1, TER119, Singlec-F, FceRIa, CD19, B220 cocktail). After staining and washing, the cells were fixed with 4% PFA and analyzed on a Becton-Dickinson FACS ARIA II, CyAn advanced digital processor (ADP). Data were analyzed using the FlowJo software. Among CD45^+^ cells, CD11b^+^ Ly6G^+^ cells were classified as neutrophils, Ly6G^−^CD11b^+^ F4/80^+^ as monocytes/macrophages, Ly6G^−^ CD11b^+^ CD115^+^ as monocytes, Ly6G^−^ CD11c^+^ MHC II^+^ as dendritic cells (DC), CD3^+^ as total T cells, CD3^+^ CD4^+^ as CD4 T cells, CD3^+^ CD8^+^ as CD8 T cells, CD19^+^ as B cells. Lin^−^ Sca^+^ cKit^+^ cells were identified as total HSCs, which were subdivided into Flt3^high^ (short-term HSC or MPP) and Flt3^low^ (long-term HSC). Lin^−^ Sca^−^ cKit^+^ cells were subdivided into CD34^low^ CD16/32^low^ (MEP), CD34^high^ CD16/32^low^ (CMP) and CD34^low^ CD16/32^high^ (GMP). The Lin^−^ CD127^+^ cKit^+^ cells were identified as CLP.

To analyze B lineage fractions, non-B cells (after lysis of red blood cells) were first dumped with FITC-CD3, -TER119, -LY6G, -LY6C, -CD11b, and -NK1.1. The remaining cells were then sequentially gated on BV650 anti-B220 (Cat. # 103241, clone RA3-6B2), APC anti-CD43 (Cat. # 143207, clone S11), PerCP-Cy5 anti-CD24 (Cat. # 101824, clone M1/69), PE anti-BP1/CD249/Ly51 (Cat. # 108307, clone 6C3), BV421 anti-IgM (Cat. # 406517, clone RMM-1), APC-Cy7 anti-IgD (Cat. # 405715, clone 11-26c.2a), PE-Cy7 anti-CD93 (Cat. # 136505, clone RAA4.1), PE-Dazzle 594 anti-CD19 (Cat. # 115553, clone 6D5).

### Single cell RNA sequencing (scRNA-seq)

Bone marrows were pooled from three mice/genotype, and red blood cells were lysed. HSCs/progenitors and B subsets (excluding mature B) were sorted as described above. In order to obtain an even coverage of each cell compartment/subset, we mixed approximately 1/6 of pre-B, ¼ of immature B (which are much more abundant than the others are) with all the HSCs/progenitors (including B progenitors). About 5×10^4^ live cells were subjected to a droplet-based 10x Genomics chromium single cell RNA-Seq on a NovaSeq 6000 sequencer, and analyzed with a Cell Ranger pipeline. Approximately 11,042 *Ubxn3b*^+/+^ and 9,269 *Ubxn3b*^−/−^ cells were sequenced and; ~1600 gene/cell and 3800 unique molecular identifier (UMI) counts/cell were obtained. Clustering cells, annotating cell clusters and analyzing differentially expressed genes were performed with Loupe Browser 5.0.1. For annotating cell clusters, we did not merely rely on the surface markers for flow cytometry, rather we referred to a database, Bloodspot, which provides transcript expression profiles of genes and gene signatures in healthy and malignant hematopoiesis and includes data from both humans and mice ^42^. The B subsets were readily identified by a medium to high level of B-restricted transcription factors (*Pax5*, *Ebf1*) and surface markers (*Cd19*, *Cd79a*), while a barely detectable level of *Fcgr3* and *Cebpa*. GMPs were *Cd19*^−^, *Cd79a*^−^, *Fcgr3^hi^*, *Cd34^h^*, *Kit^in^*, *Ly6a* (*Sca-1*)^−^, *Flt3*^−^, *Il7r*^−^, *Cebpa^hi^*. CMPs are *Cd19*^−^, *Cd79A*^−^, *Fcgr3^lo^*, *Cd34^hi^*, *Kit^hi^*, *Ly6a* (*Sca-1*)^−^, *Flt3^In^*, *Il7r^Lo^*, *Itga2b* (*Cd41*)^*Lo*^, *Cebpa^hi^*. MEPs are *Cd19*^−^, *Cd79a*^−^, *Fcgr3*^−^, *Cd34^in^*, *Kit^hi^*, *Ly6a* (*Sca-1*)^−^, *Flt3*^−^, *Il7r* ^−^, *Itga2b* (*Cd41*) ^*hi*^, *Slamf1* (*Cd150*) ^*in*^, *Cebpa^lo^ Gata1*^*hi*.^ HSCs are *Cd19*^−^, *Cd79a*^−^, *Fcgr3*^−^, *Cd34^hi^*, *Kit*^hi^, *Ly6a* (*Sca-1*) ^*in*^, *Flt3^in^*, *Il7r* ^−^. Bioinformatics analyses were performed using Reactome (https://reactome.org/). Only those genes with an average count of greater than one per cell and a p<0.1 were analyzed.

### Calcium Flux Assay

Bone marrow B factions were sorted and calcium influx was assayed with a Fluo-4 Direct^™^ Calcium Assay Kit (ThermoFisher cat# F10471). Briefly cells were incubated with Fluo-4 Direct^™^ at 37°C for 1 hour. The cells were stimulated with 20μg/ml of a purified F(ab’)2 goat anti-mouse IgM (μ chain) antibody (BioLegend, Clone Poly21571, Cat# 157102) at 37°C for 10 seconds, and were immediately analyzed by flow cytometry on a Becton-Dickinson FACS ARIA II, CyAn advanced digital processor (ADP). The final results were presented as the ratio of the mean fluorescence intensity at any given time point [F] subtracted by the fluorescence at time point zero [F0] (before stimulation) and divided by F0, i.e., ΔF/F0.

### Multiplex Enzyme-Linked ImmunoSorbent Assay (ELISA)

We used a LEGENDPlex (BioLegend, San Diego, CA 92121, USA) bead-based immunoassay to quantify the cytokine concentrations in the sera of SARS-CoV-2 infected mice. The procedures were exactly same as described in the product manual. Briefly, the samples were mixed with antibody-coated microbeads in a filter-bottom microplate, and incubated at room temperature for 2hrs with vigorous shaking at 500 rpm. After removal of unbound analytes and two washes, 25 μL of detection antibody was added to each well, and the plate was incubated at room temperature for 1hr with vigorous shaking at 500 rpm. Twenty-five μL of SA-PE reagent was then added directly to each well, and the plate was incubated at room temperature for 30min with vigorous shaking at 500 rpm. The beads were washed twice with wash buffer, and then transferred to a microfuge tube. The beads were fixed with 4% PFA for 15min and resuspended in an assay buffer. The beads were run through a BIORAD ZE5 and the concentrations of analytes were calculated with the standards included using a LEGENDPlex software.

### Quantification of serum IgG by Enzyme-Linked ImmunoSorbent Assay (ELISA)

Anti-SARS-CoV-2 Spike IgG titers were measured with a commercial ELISA kit (Acro Biosystems, Cat # RAS-T018). For quantification of influenza IgG, one nanogram of recombinant A/PR/8/34 influenza NP (generated by UConn Health Protein Expression Core) in 100 μL of coating buffer (0.05 M carbonate-bicarbonate, pH 9.6) was coated to a 96-well microplate at 4°C overnight. The plate was washed once with a wash solution (50 mM Tris, 0.14 M NaCl, 0.05% Tween 20, pH 8.0), and blocked with 4% bovine serum albumin at room temperature for 2hrs. 100μL of each diluted serum specimens (500-fold) was added a well and incubated at room temperature for 1hr, then the unbound serum was washed off three times with the wash solution. 100μL of diluted horseradish peroxidase-conjugated goat anti-mouse IgG was added to each well and incubated at room temperature for 1hr. After stringency wash, 100 μL of substrate 3,3’,5,5’ - tetramethylbenzidine (TMB) was added to each well and incubated at room temperature for 5-30min for color development, and terminated by 100 μL of 0.16M sulfuric acid. The absorption at wavelength 450nm (A_450nm_) was read on a Cytation 1 plate reader (BioTek, Winooski, VT, USA).

### Immunoblotting

To prepare B cell fractions for immunoblotting, bone marrow B cells were sorted by flow cytometry as described above (~10^4^-10^5^ cells each fraction), pelleted down by brief centrifugation, suspended in 50 μL of 2xSDS-PAGE sample buffer, boiled at 95°C for 5 min, and centrifuged at 13,000xg for 10min. Immunoblotting was performed using standard procedures. Briefly, protein samples were resolved by SDS-PAGE (sodium dodecyl sulfate-polyacrylamide gel electrophoresis, 4-20% gradient) and transferred to a nitrocellulose membrane. The membrane was blocked in 5% fat-free milk at room temperature for one hour, incubated with a primary antibody over night at 4°C, washed briefly and incubated with an HRP-conjugated secondary antibody for 1 hour at room temperature. An ultra-sensitive or regular enhanced chemiluminescence (ECL) substrate was used for detection (ThermoFisher, Cat# 34095, 32106). For immunoblotting of proteins from an extremely low number of sorted bone marrow B cells, a Lumigen ECL substrate ought to be used (Southfield, Michigan 48033, USA).

### Reverse Transcription and Quantitative (q) PCR

Up to 1□×□10^6^ cells or 10mg, tissues were collected in 350□μl of RLT buffer (QIAGEN RNeasy Mini Kit). RNA was extracted following the QIAGEN RNeasy manufacturer’s instructions. Reverse transcription of RNA into complementary DNA (cDNA) was performed using the BIORAD iScript^™^ cDNA Synthesis Kit. Quantitative PCR (qPCR) was performed with gene-specific primers and SYBR Green PCR master mix. Results were calculated using the –ΔΔCt method and a housekeeping gene, beta actin, as an internal control. The qPCR primers and probes for immune genes were reported in our previous studies ^15, 27,43^. The new primers are listed in Table 1.

### Data Acquisition and Statistical Analysis

The sample size chosen for our animal experiments in this study was estimated according to our prior experience in similar sets of experiments and power analysis calculations (http://isogenic.info/html/power_analysis.html). All animal results were included and no method of randomization was applied. All data were analyzed with a GraphPad Prism software by non-parametric Mann-Whitney test or two-tailed Student’s *t*-test depending on the data distribution. The survival curves were analyzed by a Log-Rank test. P values of ≤0.05 were considered statistically significant.

## Footnotes

### Author contribution

T.G designed and performed the majority of the experimental procedures and data analyses. D.Y. helped T.G. with most of the experimental procedures. T.L. and A.G.H. contributed to some of the experimental procedures. B.W. contributed to flow cytometry analysis. B.T. and L.H. helped to acquire the influenza data. K.W provided guidance to bone marrow transplantation experiments. Y.W., L.Y., G.C., L.H., A.T.V. and E.F. contributed to discussion, data interpretations and/or helped to improve writing. P.W. conceived and oversaw the study. T.G. and P.W. wrote the paper and all the authors reviewed and/or modified the manuscript.

### Funding Source

This project was funded in part by National Institutes of Health grants to P. W, R01AI132526 and R21AI155820, and an UConn Health startup fund to P.W.

### Conflict of Interest

No financial or non-financial interest to disclose.

### Data availability

All data generated or analysed during this study are included in this published article (and its supplementary information files). The raw RNAseq data is available at https://www.ncbi.nlm.nih.gov/geo/, GEO # (pending)

### Biological materials

All unique materials used are readily available from the authors. However, the availability of live animals may change over time.

## Supplemental Materials

Supplemental Figs. 1-9.

Supplemental Table 1-4.

Supplemental Movie 1: SARS-CoV-2-infected *Ubxn3b*^+/+^ mice on day 2 after infection. Supplemental Movie 2: SARS-CoV-2-infected *Ubxn3b*^−/−^ mice on day 2 after infection.

## Notes

### Competing Interest Statement

The authors have declared no competing interest.

### Summary of Updates

Fig.6 legend was corrected.

