## Supplemental Figs 1-6 for "An Essential Role of UBXN3B in B Lymphopoiesis"

Tingting Geng et al.

This file contains 9 supplemental figures and legends.

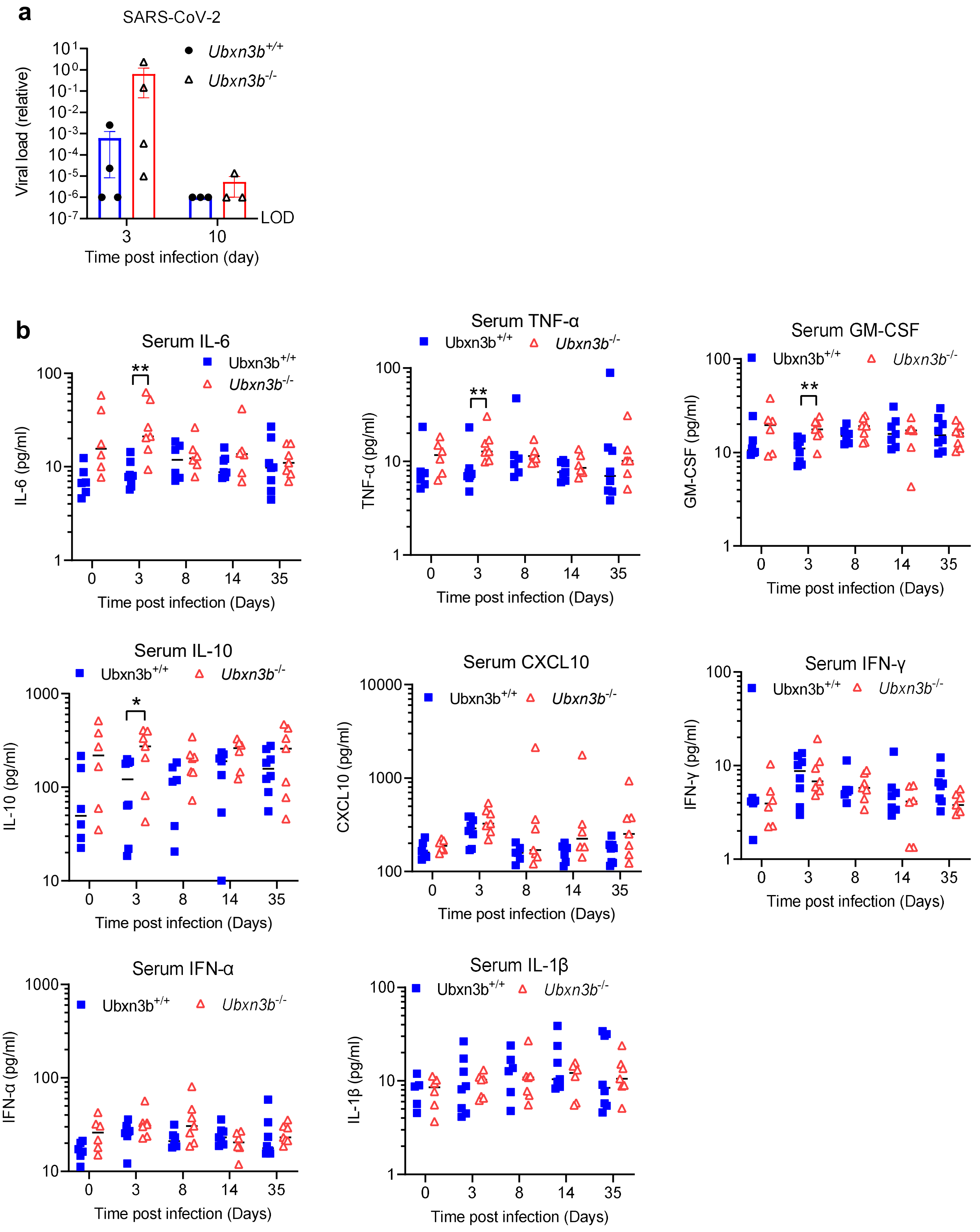


**Supplemental Fig.s1 UBXN3B is essential for controlling SARS-CoV-2 pathogenesis**. Sex-and-age matched littermates were administered $2x10^5$ plaque forming units (PFU) of SARS-CoV-2 intranasally. **a**) Quantitative RT-PCR (qPCR) quantification of SARS-CoV-2 loads in the lung at days 3 and 10 post infection (p.i). Each symbol= one mouse, the small horizontal line: the median of the result. *, p<0.05; **, p<0.01, ***, p<0.001 (non-parametric Mann-Whitney test) between *Ubxn3b*$^{+/+}$ and *Ubxn3b*$^{-/-}$ littermates at each time point.

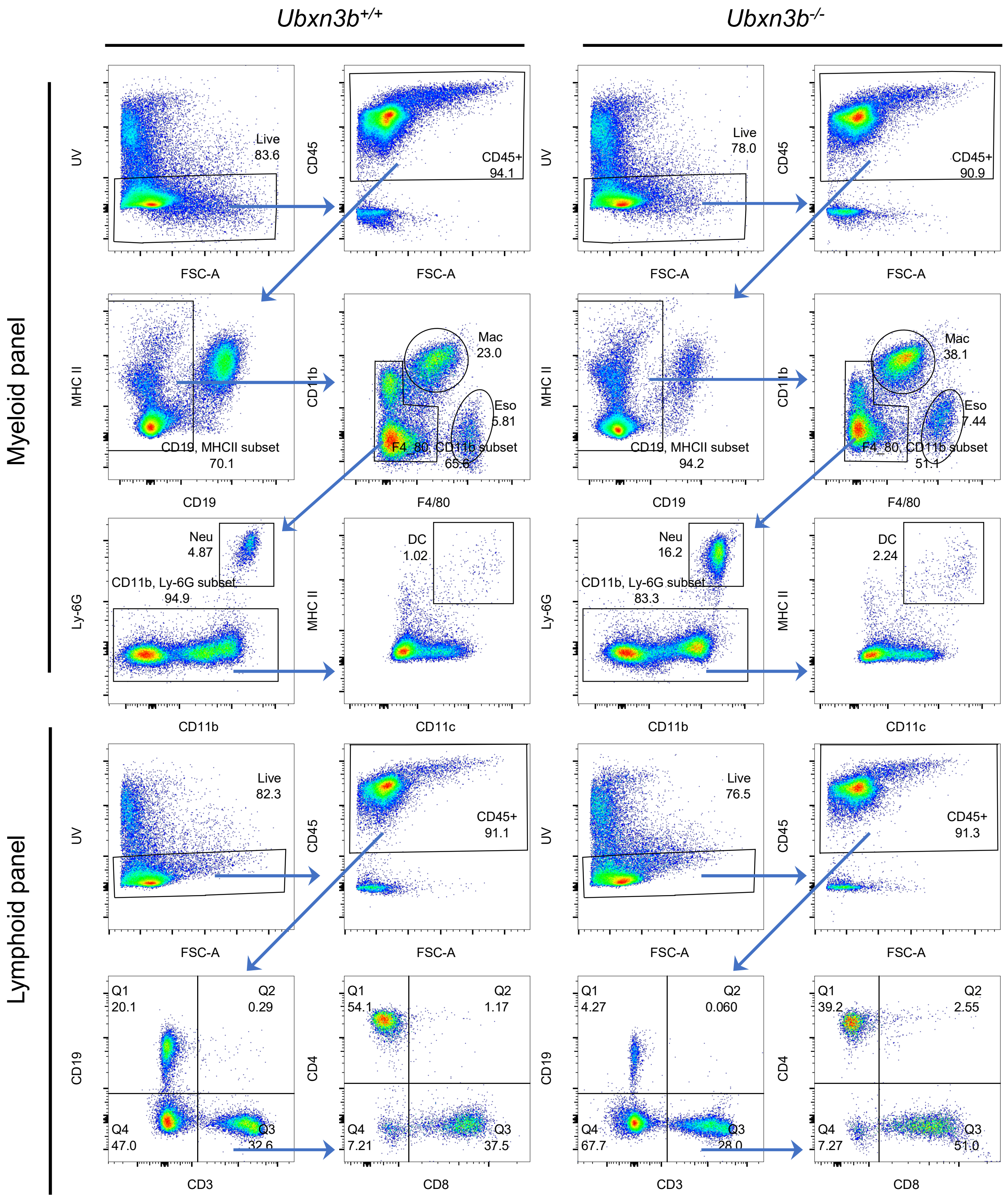


**Supplemental Fig.s2 Dysregulated immune compartmentalization in *Ubxn3b*$^{-/-}$ lung.** The graphs illustrate the gating strategy of different immune populations in the lung at day 35 post SARS-CoV-2 infection (related to **Fig.2f,g**). Mac: macrophage, Neu: neutrophil, DC: dendritic cell, Eso: eosinophil. CD3$^{+}$ T, CD8$^{+}$ T, CD4$^{+}$ T, CD19$^{+}$ B.

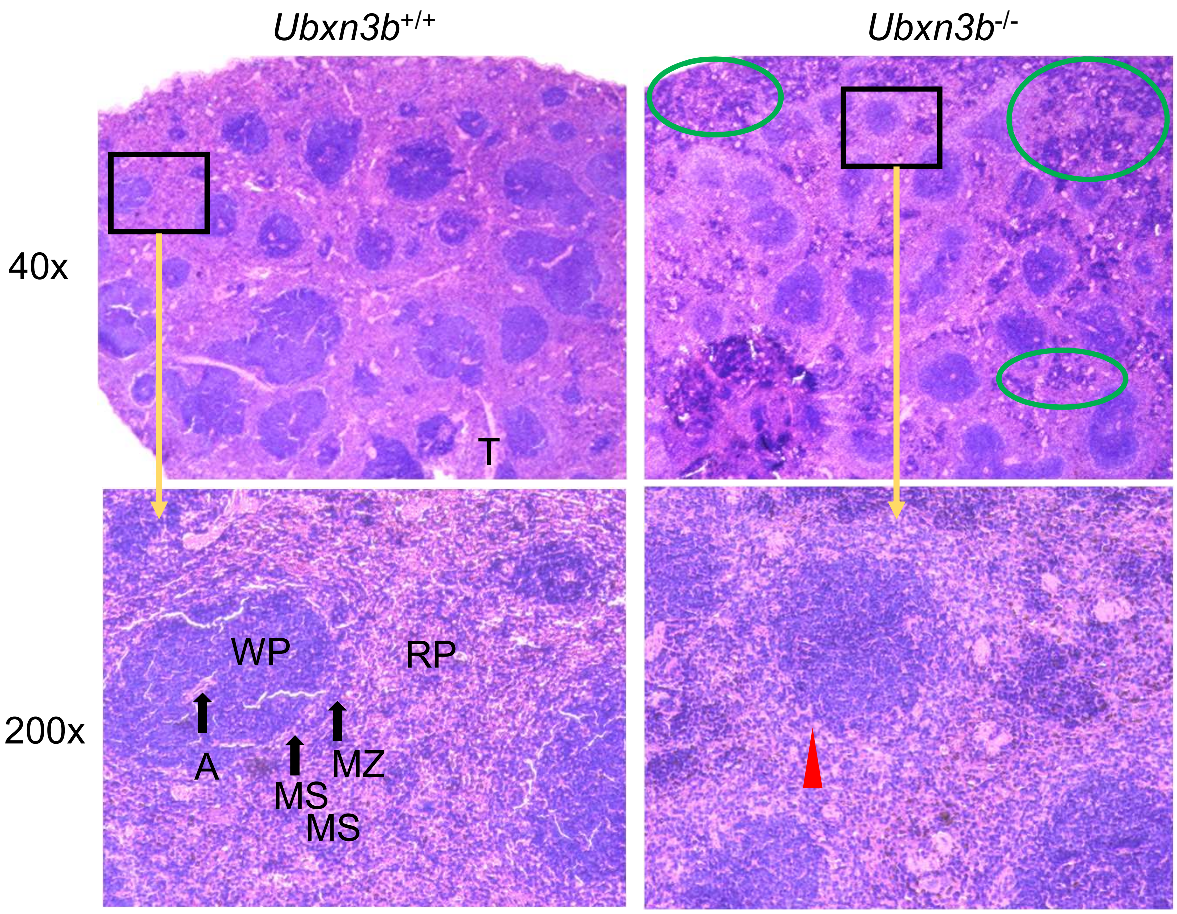


**Supplemental Fig.s3 Splenic atrophy in *Ubxn3b*$^{-/-}$ mice**. The figures show H&E staining of spleens on day 35 after SARS-CoV-2 infection. The red arrow head indicates reduced cell density in the marginal zones of white pulps. The green circles likely show the myeloid clumps in red pulps. WP: white pulp, RP: red pulp, A: central arteriole, MS: marginal sinus, MZ: marginal zone, T: trabecula.

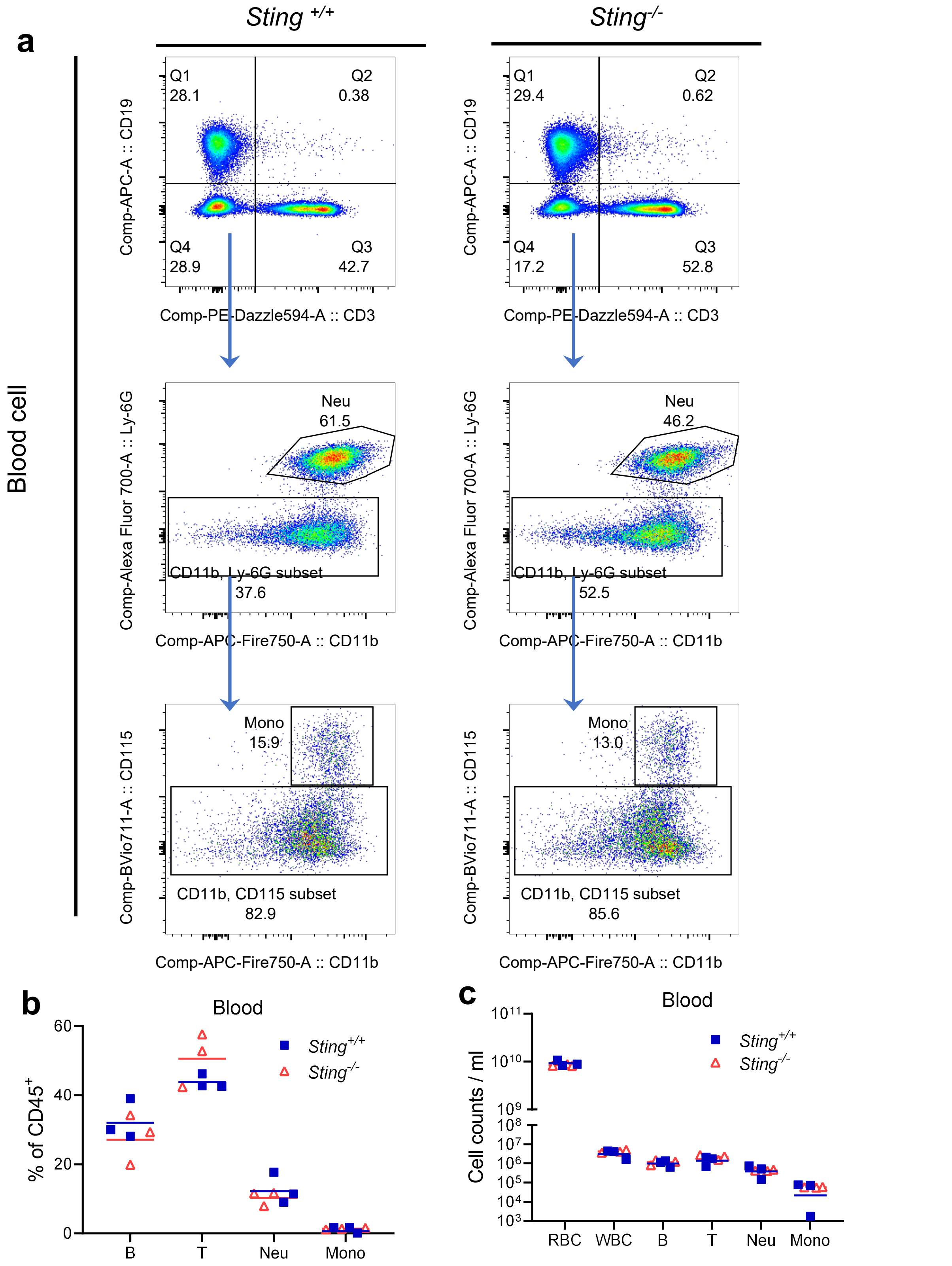


**Supplemental Fig.s4 STING is dispensable for steady-state hematopoietic homeostasis**. **a**) The gating strategy, **b**) The percentage and **c**) counts of each blood cell type in *Sting*$^{+/+}$ and *Sting*$^{-/-}$ littermates, quantitated by flow cytometry. Each symbol=one animal. The horizontal line indicate the mean of the results. RBC: red blood cell, WBC: white blood cell, CD19$^{+}$: B cell, CD3$^{+}$ : T cell, Neu: neutrophil, Mono: monocyte.

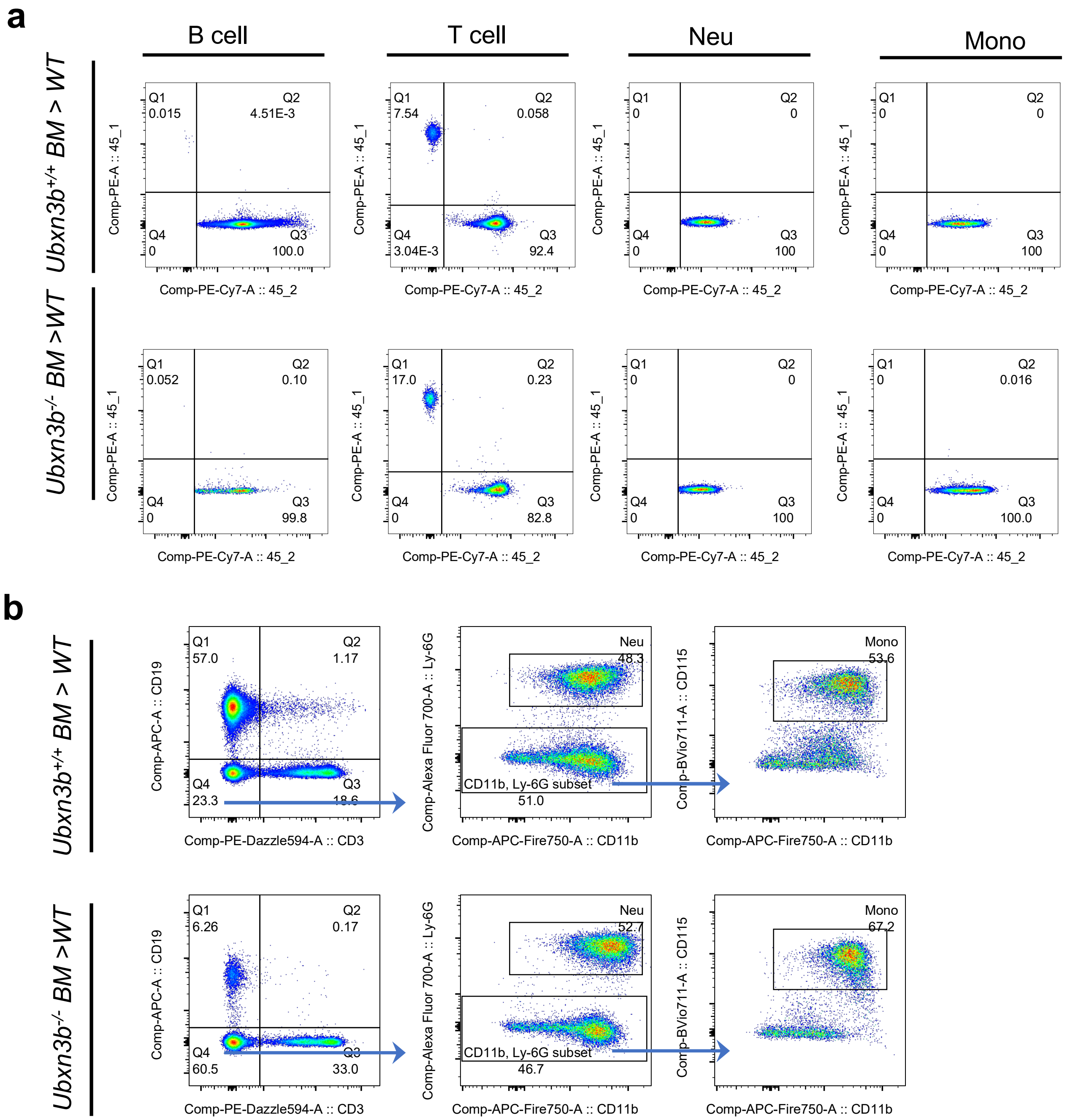


**Supplemental Fig.s5**. **UBXN3B plays a cell-intrinsic role in B lymphopoiesis.** Irradiated wild type (WT, CD45.1 ) mice were transplanted with *Cre*$^+$*Ubxn3b*$^{f/f}$ (CD45.2) bone marrow . The mice were then treated with tamoxifen (TMX) to delete Ubxn3b (designated *Ubxn3b*$^{-/-}$ BM–WT) or not (designated *Ubxn3b*$^{+/+}$ BM–WT) in hematopoietic cells. **a**) The percentage of blood CD45.1 and CD45.2 cells 1.5 month after bone marrow transplantation (BMT). Over 99% B/Neu/Mono are CD45.2$^+$, over 82% T cell are CD45.2$^+$, indicating successful irradiation and reconstitution. **b**) The gating strategy of blood immune cells in BMT mice (related to **Fig.4**). B: CD19 $^+$ B cell, T : CD3$^+$ T cell, Neu: neutrophil, Mono: monocyte.

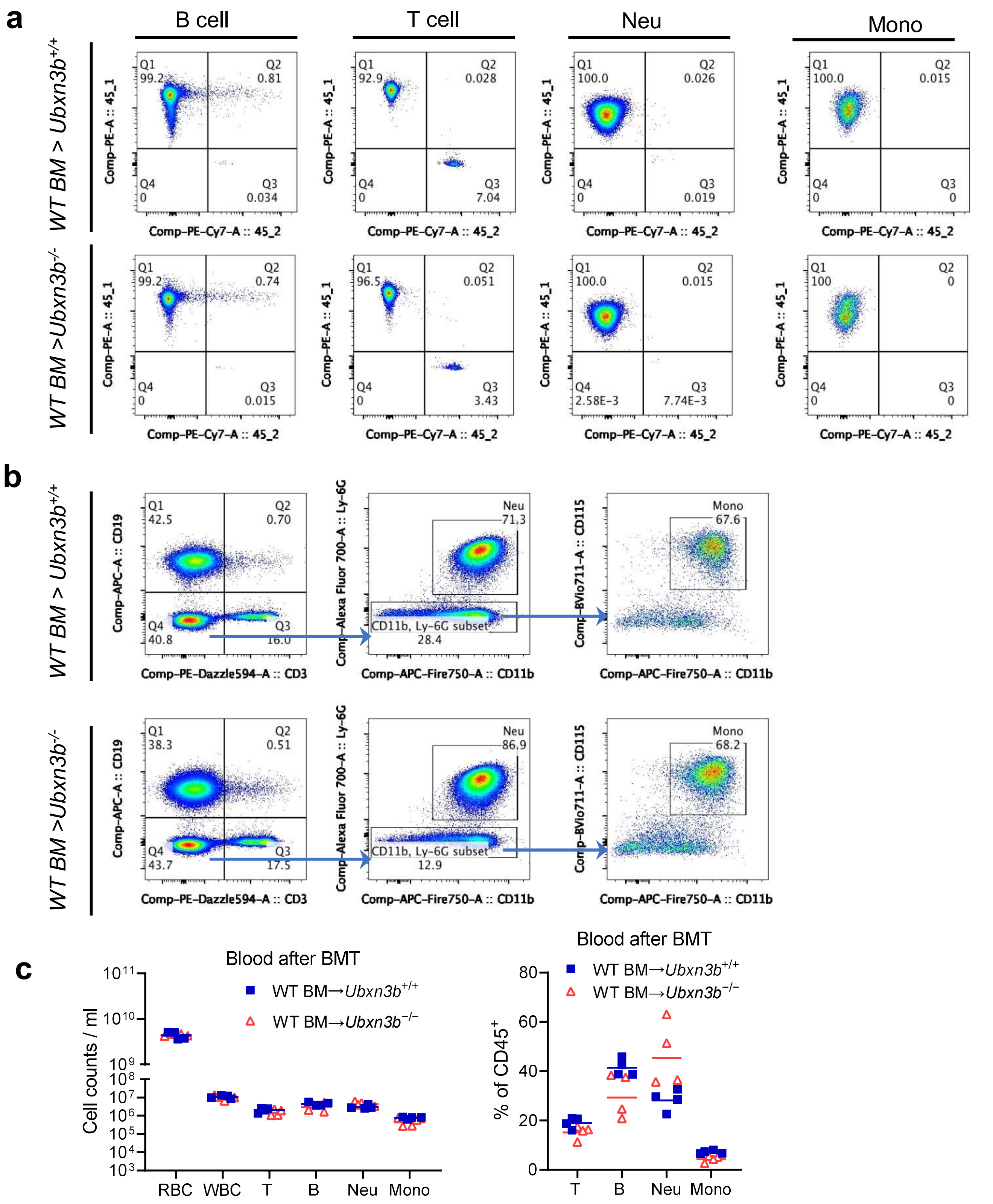


**Supplemental Fig.s6**. **UBXN3B plays a cell-intrinsic role in B lymphopoiesis.** Irradiated *Cre*$^{+}$*Ubxn3b*$^{f/f}$ and *Ubxn3b*$^{f/f}$ (CD45.2) mice were transplanted with wild type (WT, CD45.1) bone marrow . The mice were then treated with tamoxifen (TMX) to delete Ubxn3b (designated WT BM–*Ubxn3b*$^{-/-}$) or not (designated WT BM–*Ubxn3b*$^{+/+}$) in non-hematopoietic cells. **a**) The blood CD45.1 and CD45.2 cells 1.5 month after bone marrow transplantation (BMT). Over 99% B/Neu/Mono are CD45.1$^{+}$, over 92% T cell are CD45.1$^{+}$, indicating successful irradiation and reconstitution. **b**) The gating strategy of blood immune cells in BMT mice. **c**) The count and percentage (relative to CD45$^{+}$) of each blood cell type in BMT mice at 15 days after induction of Ubxn3b deletion by tamoxifen (TMX), quantitated by flow cytometry. *, P<0.05; **, P<0.01, ***, P<0.001 (two-tailed Student's *t*-test). Each symbol=one animal. The horizontal line indicate the mean of the results. RBC: red blood cell, WBC: white blood cell, B: B cell, T : T cell, Neu: neutrophil, Mono: monocyte.

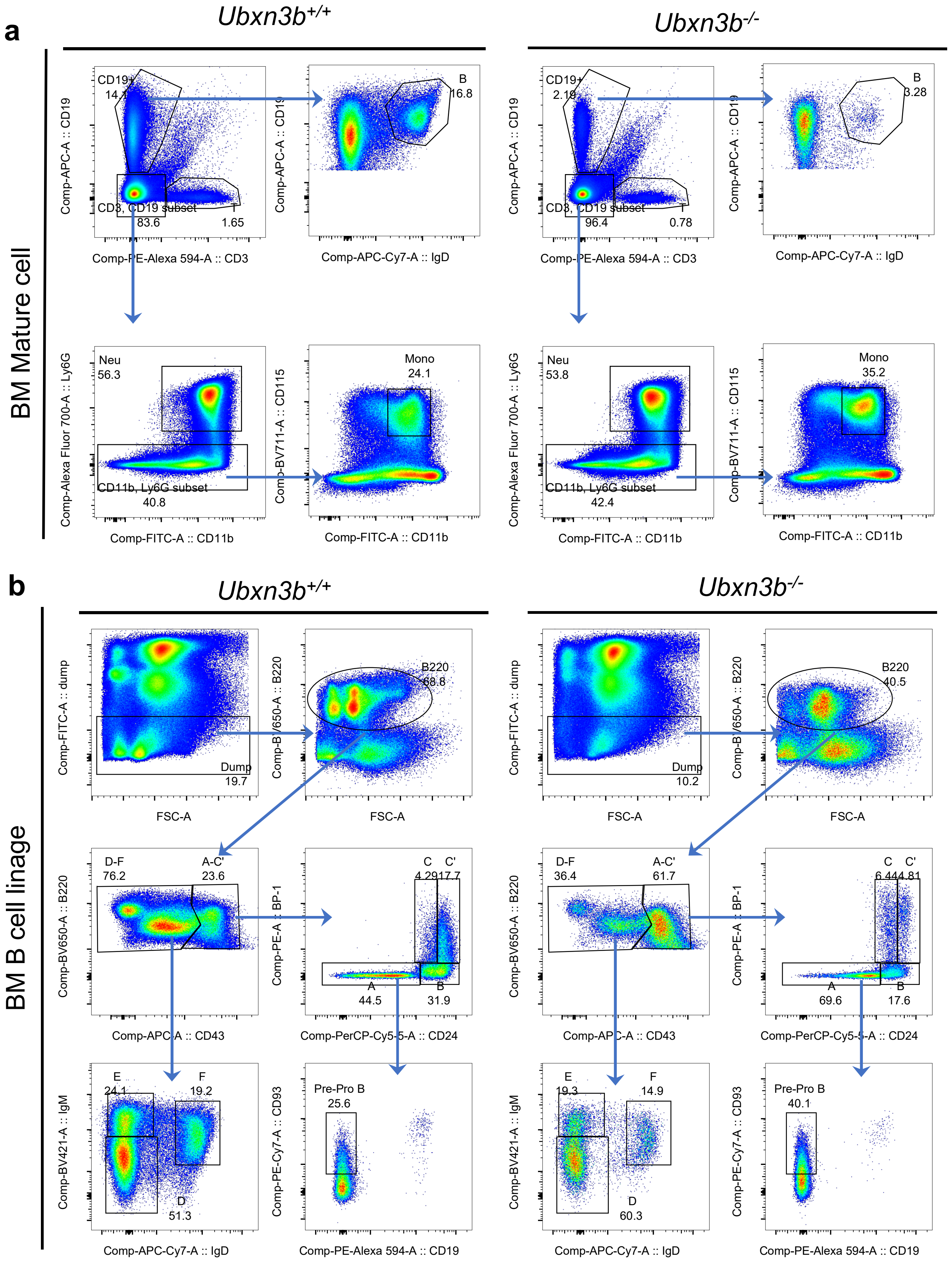


**Supplemental Fig.s7 UBXN3B is essential for early B lymphopoiesis in the bone marrow**. The gating strategy for **a**) terminally differentiated (related to **Fig.5a**) and **b**) B lineage subsets in the bone marrow of *Ubxn3b*$^{+/+}$ and *Ubxn3b*$^{-/-}$ littermates (related to **Fig.5b**). Fraction A: Pre-pro-B ( pre-progenitor B), Fraction B: pro-B (progenitor B), Fraction C: pre-BI, Fraction C': large pre-BII, Fraction D: small pre-BII, Fraction E: immature B, Fraction F: recirculating mature B. T : T cell, Neu: neutrophil, Mono: monocyte.

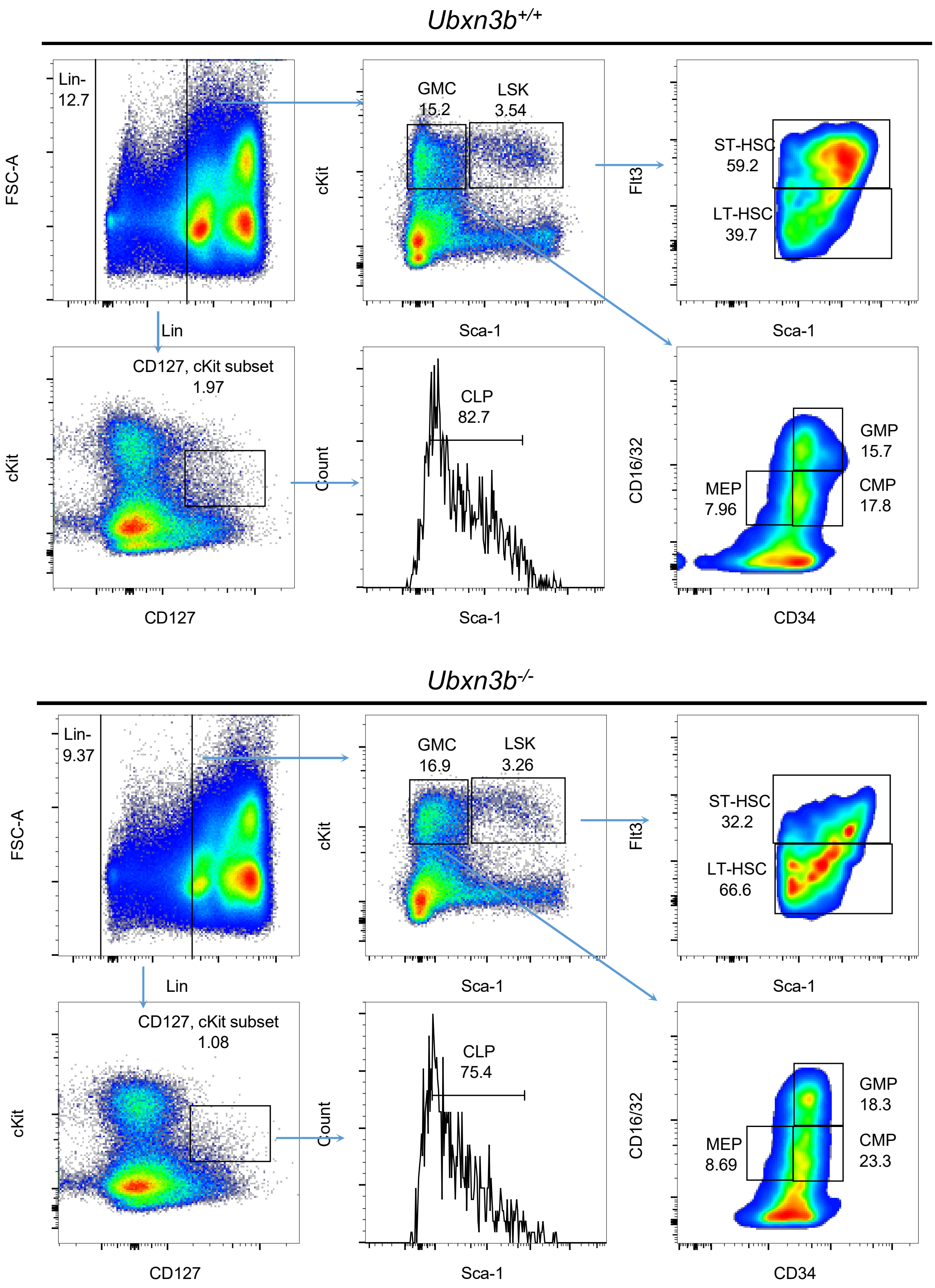


**Supplemental Fig.s8 Bone marrow stem cells and CLPs are modestly reduced in *Ubxn3b*$^{-/-}$ mice.** The gating strategy for hematopoietic stem cells (HSC, Lin$^{-}$, Sca1$^{+}$ Kit$^{+}$) and lineage progenitors (related to **Fig.5c**) in the bone marrow of *Ubxn3b*$^{+/+}$ and *Ubxn3b*$^{-/-}$ littermates. ST-HSC; short-term HSC, LT-HSC: long-term HSC, CLP: common lymphoid progenitor, CMP: common myeloid progenitor, GMP: granulocyte-macrophage progenitor, MEP: megakaryocyte-erythrocyte progenitors.

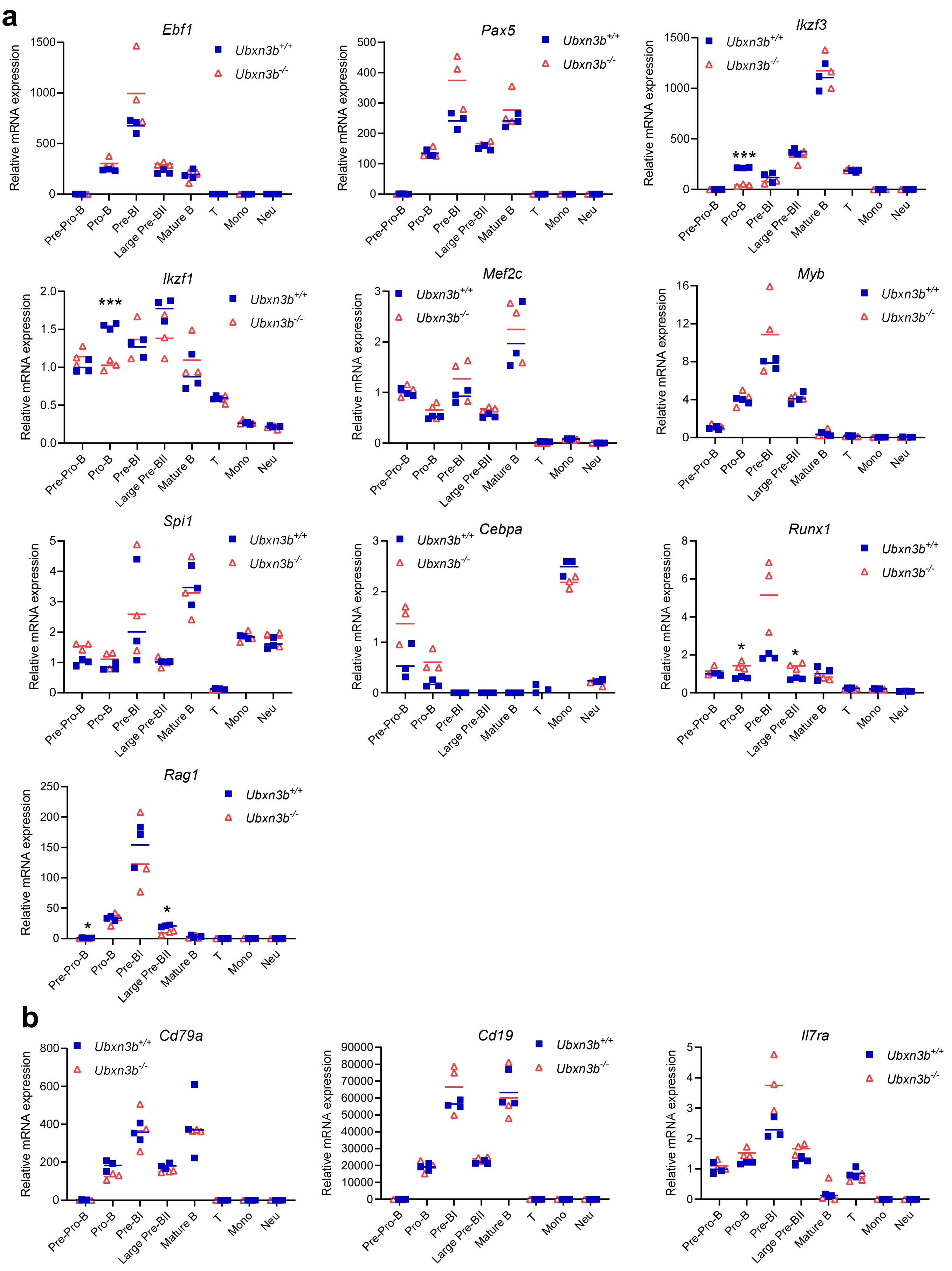


**Supplemental Fig. s9 qRT-PCR quantification of transcription factors and surface markers** in steady-state *Ubxn3b*$^{+/+}$ and *Ubxn3b*$^{-/-}$ littermates. The bone marrow cell compartments were sorted by flow cytometry and the mRNA expression of **a**) transcription factors and **b**) cell surface markers was quantitated by qRT-PCR and normalized to a house keeping gene, *Actb*. Each symbol=one animal. The horizontal line indicates the mean of the results. *, P<0.05; **, P<0.01, ***, P<0.001 (two-tailed Student's *t*-test).
